# Simulating the dynamics of gene regulatory systems at cellular scale

**DOI:** 10.64898/2026.11.01.763459

**Authors:** Lukas Maischak, David Jordan, Stefan Manolache, Hajk-Georg Drost

**Author notes:** Authors contributed equally (Lukas Maischak, David Jordan, Stefan Manolache).

## Abstract

The temporal nature of eukaryotic gene expression is the result of a complex, non-linear, dynamical system of regulatory processes. So far, models of such systems trade realism for tractability by using strict simplifying assumptions. These simplified models are often unfaithful to our biological understanding; however, without such simplifications, analytical solutions become more challenging and these dynamical systems must then be studied through simulation. Here, we explore the question of how this complex web of gene regulatory connections can be faithfully simulated for all tens of thousands of genes that usually define a eukaryotic organism. We present a proof-of-principle model and simulator, *GeneRegulatorySystems.jl* which is based on a biophysical chemical master equation that overcomes three key restrictions. First, it allows an exact stochastic simulation of fully non-linear gene expression dynamics, including feedback control, to parsimoniously capture cause and effect relationships of gene regulation. Further, it includes a general framework for implementing diverse mechanisms of gene control such as protein mediated degradation, RNAi, and protein complex formation. Finally, it scales realistically to tens of thousands of genes. We achieve this by introducing a domain-specific programming language to aid the design of experimental protocols from small-scale, well-studied, scenarios of gene regulation inspired by dynamical systems biology up to large-scale, single cell omics experiments with interventions. In this way, our framework provides a grammar of *in silico* experimental design that can capture single-cell gene expression dynamics for entire genomes.

## Introduction

Despite the incredible amount of phenotypic variation found on our planet, the processes that govern development as a reproducible procedure to create species-characteristic phenotypes are remarkably robust. Although much has been learned about individual genes, transcription factors, or biological pathways that are involved in the regulation of various developmental processes (*1*), it is not clear why some stages of development are more prone to perturbations than others (*2*). These susceptible periods of developmental organisation are often the causal origin of many developmental diseases, because their overall systemic instability introduces opportunities for significant change that often result in deleterious effects (diseases) but can sometimes also yield innovative outcomes (heterochrony, adaptation, and speciation) (*3* – *5*).

Understanding development requires a systems biology perspective, as many different molecular and morphological scales are involved in governing the developmental milestones that ultimately result in the formation of a healthy body (*6* , *7*). Such insights promise not only a better understanding of how developmental programs are evolved to sufficiently reproduce complex body structures across generations, but also reveal the key genes responsible for many known and yet incurable diseases. Here, we approach the systems biology of developmental regulation by accurately modeling the biophysics and kinetics of genome-wide cellular-scale gene regulation. Models of gene regulation face a trade-off between biophysical detail and scalability (*8*). Mechanistic descriptions of transcription, processing, translation and their regulation are usually built for a handful of genes, but each additional gene introduces a new set of rate constants. When such models are extended to hundreds or thousands of genes, the number of parameters grows rapidly. In this case, simulation becomes computationally expensive, and the parameters can no longer be constrained by the available data, so fitting becomes ill-posed and prone to overfitting (*9* –*12*).

Consequently, such methods have introduced simplifying heuristics when scaling to cell-level gene regulatory models, which result in a coarse understanding of systemic regulation.

In addition, many classical systems biology approaches focus on individual interactions and thus fail to take advantage of the interaction structure recurring throughout the network. This could be addressed through better tooling and model construction from standard templates.

In general, a complementary approach would be to use the limited available data more economically through less general, more constrained models. However, if these constraints are not chosen faithfully, the resulting models are unlikely to capture the nature of gene expression accurately enough. A promising source for parsimonious constraints are biophysical models based on the known molecular interactions that underlie gene regulation. Previously, such assumptions have often been understood to result in models that are too detailed to capture at scale because they are too complex to treat analytically or too expensive to simulate sufficiently. This has led to largely distinct approaches from the fields of single molecule biophysics, systems and synthetic biology, and experimental “omics” disciplines.

In single-molecule biophysics, experimenters tend to prefer highly accurate quantification of one or a few molecular species, using techniques such as single-molecular RNA FISH (*13*) or MS2 based localization *in vivo* (*14*), with a focus on the statistical features representing gene expression variability. In systems biology, the aim has been to capture the dynamics of small gene regulatory motifs with analytical treatments, often based on mathematical tools and methods from dynamical systems theory. These include phase portraits, the study of attractors and stability, and bifurcation analysis (see e.g. (*15*)). Finally, in large-scale omics and bioinformatics approaches based on nucleic acid sequencing (*16* , *17*), the biophysical details and dynamics of gene regulation are often ignored in favor of associational methods based on gene expression snapshots which are assumed to be representative of some quasi-stable steady state.

These are often necessary assumptions because sequencing based methods are not well suited for observing state over time, as they are invasive procedures and often require isolating and destroying the measured cell. And while these methods allow thousands of molecules to be measured at once, each estimate of an individual gene’s expression is more noisy (*18*). Thus, the ideal approach would explicitly distinguish classical experimental designs based on extracting whole cells from an organism to sample RNA libraries, and others where cells are kept alive, e.g. using fluorescence readouts of mRNA and proteins (*19* , *20*) to non-invasively monitor dynamics. New technologies such as LIVE-Seq (*21*) can bridge this gap by capturing RNA molecules from the same living cell over time.

Thus far it has not been possible to integrate the diverse perspectives and the technical methods of these fields into a single quantitative modeling framework. Ideally, any such unified framework should allow the generation of physically realistic models at the microscopic level but should also represent abstractions at network-level scales. Constraints need to be chosen so that candidate models are simple enough to reason about effectively but, at the same time, permit complex dynamics to arise purely from endogenous gene regulatory interactions. Finally, any such model should be falsifiable by permitting arbitrary exogenous interventions and as a result enabling the generation of simulated data that adheres as closely as possible to the structure of real omics experiments. Overall, a useful toolkit must incorporate the diverse theoretical perspectives and match the predictions of each of these fields at their respective scales: from the expression of single genes, through the regulation within small biological pathways, and ultimately up to the large scale gene expression trajectories of entire cells.

## Results

In studies using biophysical models of gene regulation, analytic treatments such as those employing generating functions (*22*) or moment closure methods (*23* –*25*) are the gold standard and the quantitative insights gained from their use have been foundational (*13*). However, applying these methods to complex networks often requires simplifying assumptions to make the analysis feasible (*26*). Analytical treatments of systems under feedback control are even more challenging (*27* , *28*). Therefore, for the complex networks of interactions found in many gene regulatory systems, simulations remain the only effective alternative to explore their behavior. There is a risk that poorly constructed simulations, rather than generating insights, would merely generate idiosyncratic and unrealistic anecdotes. However, done properly, simulations have the potential to provide a top-down complement to analytically tractable bottom-up approaches, and thus provide the embankments on which a useful bridge between the two might be constructed. *Ad hoc* modeling that does not faithfully represent the underlying biophysics of gene regulatory networks is insufficient for this purpose. But it remains to be shown how biophysically realistic models of gene expression and regulation can be assembled in practice and simulated efficiently at scale. Here, we present a proof-of-concept gene regulatory systems simulator that answers this question affirmatively.

### A. A unified simulator to model the bio-physics of gene expression for systems biology

One way to accurately model the stochastic nature of gene expression is to formulate a chemical master equation (CME) (*29*), which can then be solved or simulated. Methods for solving the CME in its full distributional form for systems of this size are not currently known; however, trajectories can be sampled from its associated continuous-time Markov jump process. In this regard, we conceptualize the system as a chemical reaction network and its associated Markov process, which account for the reaction rates of the physical processes involved in gene regulation (App. A). Our formulation includes stochastic switching of promoter states, regulation of the promoter by transcription factors such as enhancers and repressors, formation of elongation complexes, nascent mRNA transcription, splicing and pre-mRNA processing (in eukaryotes), and protein translation (Fig. 1A and App. B). Such a detailed model may permit analytical solutions using generating function methods for a single gene, but multiple genes with regulatory feedback loops must be simulated or approximated, for example, using moment closure methods to derive chemical Langevin equations (*30* , *31*). In addition to transcriptional regulatory networks that include both activating and repressing links, our model also permits an arbitrary molecular interaction network that allows the inclusion of other regulatory processes such as ubiquitin-mediated proteolysis, post-transcriptional regulation by small RNAs, and transcription factor dimer and multimerization, for example. Using these features, we have recreated a number of results from classic single molecule biophysics and from dynamical systems and synthetic biology, demonstrating the power and flexibility of this unified modeling framework.

**Figure 1:**
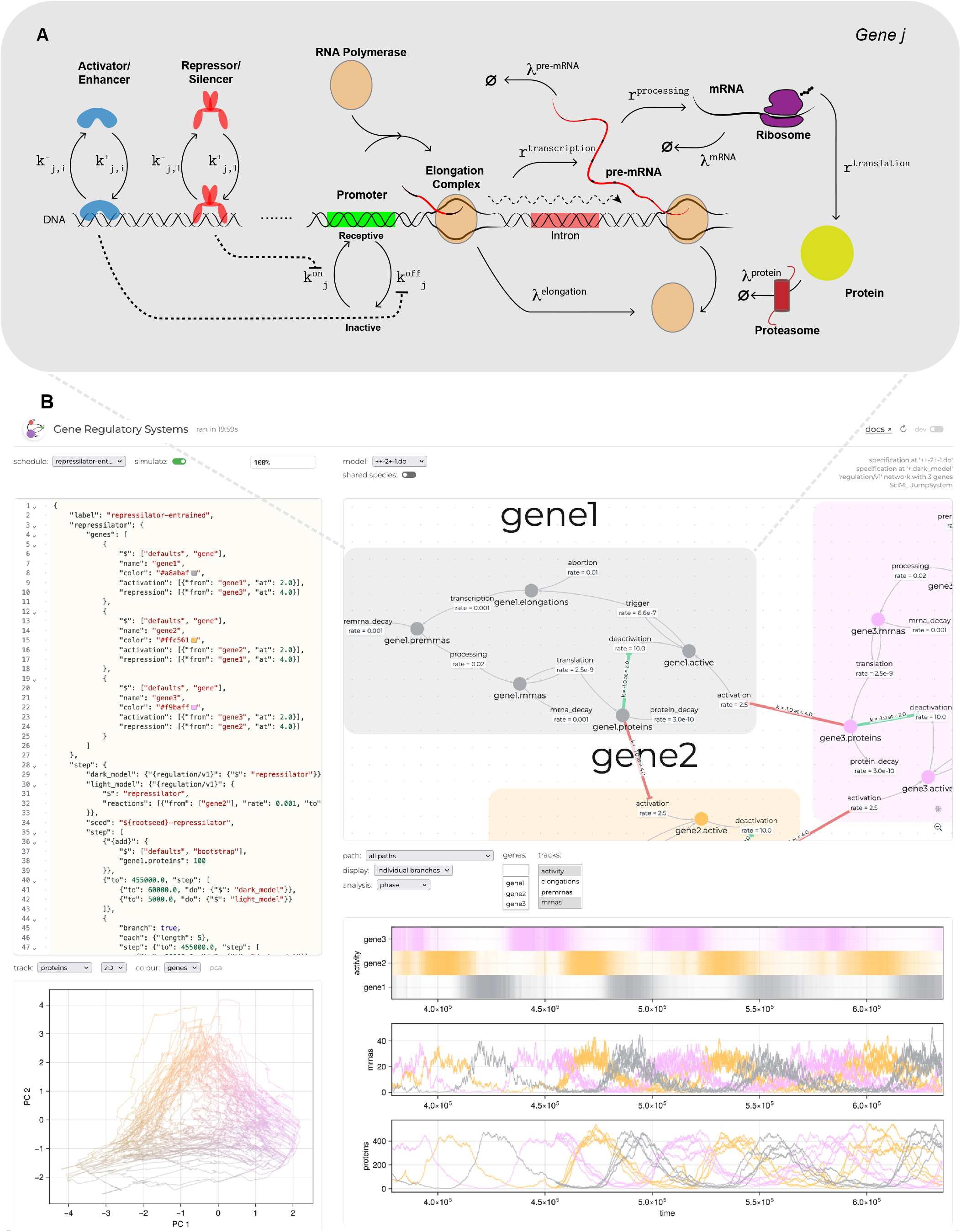
*GeneRegulatorySystems.jl* models gene expression cascades using a design language and provides an interactive user interface. (**A**) Each gene *j* is modeled as a cascade of chemical reactions with mass-action kinetics. The binary state of each promoter (by default a single copy per gene) switches with rates 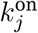 and 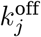 and determines whether it is receptive to the binding of RNA polymerase to form an open elongation complex. Elongations spontaneously finish to produce transcripts. The figure shows the eukaryote model where transcripts are then processed before they are available for translation by a ribosome. These reactions have rate constants 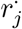. The resulting protein is then available to participate in other reactions, which may include transcriptional regulation by tempering the 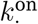 and 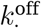 rates of the same or another gene. Besides the promoter state, the gene’s dynamic components are modeled as molecular species (elongations, nascent and mature transcripts, and proteins), all of which decay with rate constants 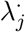. Elongation complex initiation, translation and protein degradation rates depend proportionally on the present amounts of RNA polymerases, ribosomes, and proteasomes, which are respectively modeled as molecular species but do not decay (by default). (**B**) Corresponding user interface for *GeneRegulatorySystems.jl*, highlighting how this reaction-level formalism is assembled into a modular chemical reaction network (top-right), the system dynamics it generates (bottom), and its specification, through a structured design language, which can be altered by the user (left).

Here, we present *GeneRegulatorySystems.jl*, a software package designed to construct complex models of single-cell gene regulation from modular templates and coordinate reproducible simulation through a flexible scheduling mechanism (Box 1) and output format. The basic building block of these models is a simplified “gene expression cascade” (Fig. 1A) which reflects our current understanding of the biophysics of gene regulation and expression, from promoter activation through transcription and processing, and finally to translation. These gene instances may then be assembled into larger systems (Fig. 1B) by introducing idealized regulatory connections, including feedback loops, and arbitrary additional mass-action reactions. In this way, the complexity inherent in the CME describing the reactions that comprise such a system is subsumed into a readily manipulable abstraction that nevertheless retains the detail necessary to be biophysically accurate.

Exact stochastic simulations are required to recapitulate many of the empirically observed properties of expression realistically, especially at low expression levels. Despite our simplifying assumptions, when simulated in this way, our models recapitulate important statistical features of gene expression, such as expression bursting and expression mean–variance relationships (Results B). Surprisingly, despite this complexity, these models can be efficiently simulated at scale using off the shelf algorithms, here in particular those available from the SciML ecosystem (*32*). Interoperability with these tools also introduces the possibility of additional analysis, such as an investigation of steady state behavior, or potential simulation speed ups through approximation by ordinary differential equations (ODEs) or stochastic differential equations (SDEs) (*33*).

The scheduling mechanism (Results C) enables planned interventions at arbitrary time points on both the states and the parameters, which can, for example, be used to determine the frequency response of systems to periodic inputs, or to map the dynamical phase space of smaller systems using grid searches (Results D). Most real-world experimental data at the cellular scale come from nucleic acid or protein sequencing experiments, which do not have the same time resolution as the data derived from single molecule or small network studies. In order to compare the simulations from this toolkit with real experimental data, *GeneRegulatorySystems.jl* includes a separate observation model. This model incorporates additional uncertainty introduced by the random extraction and amplification steps in sequencing protocols. The scheduling system can also be used to branch simulation trajectories such that multiple independent realizations will proceed from the system’s state at a specific time point. This can be used to generate synthetic collections of cells which share a common history, useful for sensitivity analysis and for efficiently generating pseudo single cell data.

The *GeneRegulatorySystems.jl* toolkit thus permits the generation both of exact trajectories that trace every molecular event, as well as snapshots at specific time points which can be exported as counts matrices and readily used in downstream analysis pipelines originally built for experimental data, for example with trajectory or gene regulatory network inference, denoising or imputation, and data exploration and visualization tools. This combination of realistic data with known counts, trajectories, and underlying network structures provides a unique opportunity to benchmark such analyses and, taking known intervention specifications into account, even to apply causal reasoning up to the counterfactual level.

*GeneRegulatorySystems.jl* exemplifies how to structure gene expression simulators capable of generating realistic behaviors, from single molecules to thousands of genes, including cell differentiation without the need to impose exogenous control (Results E). To accomplish this, we have leveraged state of the art simulation algorithms from DifferentialEquations.jl (*32*), in particular JumpProcesses.jl (*34*) through Catalyst.jl (*35*), and applied them to sophisticated biophysics models (*24* , *25* , *36*).

To orchestrate such simulations, we have further built a flexible yet powerful specification language, a custom scheduling system for interventions, and an observation model that mimics experimental protocols. *GeneRegulatorySystems.jl* aims to be a tool for connecting specific changes in structural assumptions with the resulting differences in gene expression dynamics, which we consider a necessary step toward a more complete understanding of gene regulation.

### B. Realistic simulations of temporal gene expression from single molecules to whole cells

The model that underlies *GeneRegulatorySystems.jl* is capable of producing realistic properties of gene expression without *ad hoc* extensions or explicit fitting. For example, the model recreates well known relationships between gene expression levels and gene expression noise that have both theoretical and experimental support (*23* , *24*). Extensive previous work has related many aspects of promoter architecture to the Fano factor, the ratio of variance in gene expression to its mean (*13* , *25*). In conditions when a gene is constitutively expressed, mRNA counts are known to be Poisson distributed. Poisson distributions are characterized by a single parameter which defines both the mean and variance simultaneously, resulting in a Fano factor of 1. Under regulation, the switching rates of the promoter are affected and the Fano factor will increase. If regulatory factors affect the on-rate rather than the off-rate, it will result in a different relationship between the Fano factor and the mean expression and the resulting curve will have a different shape (Fig. 2A,B).

**Figure 2:**
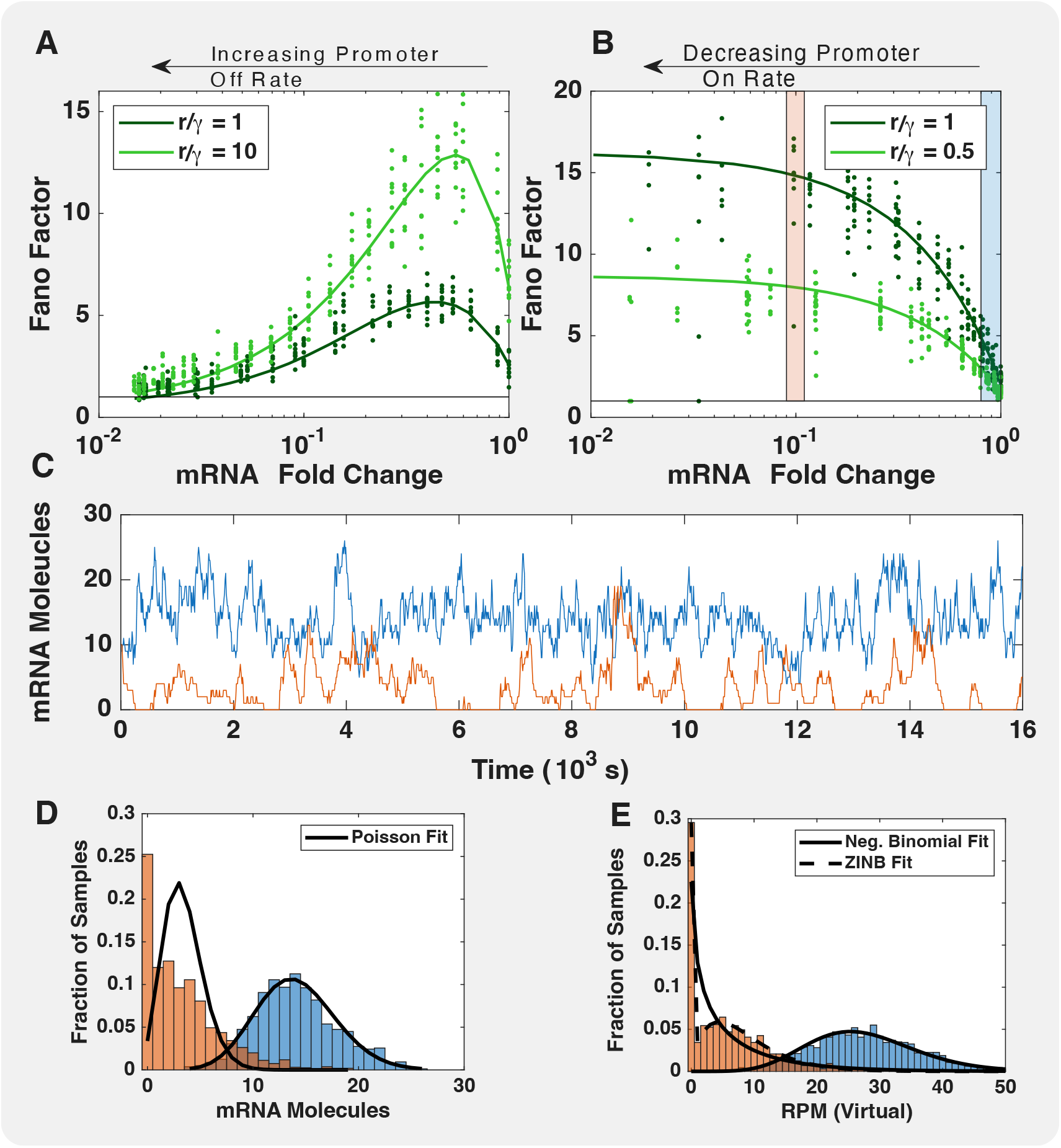
Biophysically inspired models can reproduce relevant statistics of expression at the single-gene level. *GeneRegulatorySystems.jl* can be used to to reproduce quantitative results from biophysics such as the influence of promoter architecture on gene expression noise. (**A**) and (**B**) show how changes in promoter on and off rates affect the Fano factor. For each setting, ten independent replicates are shown. The solid lines are fit-free predictions from moment-closure approximations (App. C). (**C**) Time series of the mRNA counts of two genes, one constitutively expressed (blue) and one under repression (orange) resulting in transcriptional bursting, correspond to the respective colored regions in the top right plot. The distribution of mRNAs for the constitutively expressed gene (**D**, blue) is well fit by a Poisson distribution. The bursty behavior of the repressed gene (orange) leads to zero inflation in the true counts (**D**, orange). After extraction, the read count distribution for the blue gene is well fit by a negative binomial distribution (**E**). The negative binomial fits poorly for the bursty gene (orange) as the read count distribution remains zero inflated.

Another related example is the well known fact that gene transcription tends to be “bursty” (*37*). This burstiness is believed to come from stochastic promoter switching and is often added to a model by drawing the number of molecules of mRNA produced per burst from a distribution. Although this modeling choice is essential to analytically solve models of gene expression in closed form (*26* , *38*), such an extension is not necessary to achieve bursty transcription with *GeneRegulatorySystems.jl* because the underlying promoter state explicitly switches stochastically (Fig. 2C). In our system, the burst size emerges as an effective parameter resulting from the dynamics of promoter activation and inactivation, which is influenced by other activating and repressing transcription factors. Bursty transcription is one of many biological features that can cause the so-called “zero-inflated” distributions in sampled data, such as RNA read counts (*39*). Other biological causes include temporal regulation of the gene that results in periods of no expression or cell-type-specific regulation in which the gene is expressed only optionally.

In genes consitutively expressed in a high number, the molecule counts should instead be Poisson distributed (Fig. 2D), and this behavior is recapitulated by our model. However, the read counts distributions generated by standard techniques, such as RNA sequencing, tend to be negative-binomially distributed (*36* , *40*) which we can also fit (Fig. 2E). This is often added *posthoc* as a feature of a simulation tool, but we have included a separate extraction model customized for RNA sequencing protocols (Methods J). The extraction model simulates sequencing in three stages: First, molecules are sampled at random from the pool of actual molecules in the cell without replacement. Next, molecules are selected randomly for amplification over multiple (user defined) rounds. In each round, selected molecules are copied or removed with specified probabilities that can be interpreted as the efficiency of a round of polymerase chain reaction (PCR) amplification. Finally, molecules are randomly sampled from this amplified pool to achieve a specified target molecule count. Using this synthetic extraction protocol, we naturally generate read count distributions that are fit well by negative binomial distributions for constitutively highly expressed genes (Fig. 2E, blue) and zero-inflated negative binomial distributions for lowly expressed and bursty genes (Fig. 2E, orange).

The explicit sampling procedures that we have included are also amenable to generatingfunction-based approaches to convert molecule distributions into read distributions (*22*). The stationary noise statistics in Fig. 2 are not independent of the gene’s dynamics. For a promoter switching between active and inactive states, the noise in the output is coupled to the speed with which the promoter responds to a change in its regulatory input. At a fixed metabolic cost there is an optimal trade-off between the two, and any gene with a non-vanishing leak expression will be noisier, slower, or both, than a leak-free gene of the same cost (*41*). Negative autoregulation partially relieves this constraint. For low but non-zero leak rates, an auto-repressed gene both switches on and off faster and has lower output noise than an unregulated gene of equal cost (*42*). In GeneRegulatorySystems.jl, promoter switching, the expression cascade, and feedback links are all represented explicitly, so these trade-offs follow from the kinetic parameters without needing to be imposed in an *ad hoc* manner.

*GeneRegulatorySystems.jl* can also be used to simulate large scale bulk RNA-seq datasets and to validate tools used to analyse these data, to detect differentially expressed genes, for example. We tested DESeq2 (*43*), a popular method to identify differentially expressed genes in RNA sequencing datasets. In two large systems designed to simulate *E. coli* and human datasets, gene expression levels were tuned to introduce a known number of differentially expressed genes. In addition to the single-gene properties reproduced in Fig. 2, many multi-gene statistical features of gene expression data-sets can also be faithfully simulated. In these examples, the distribution of gene expression means (Fig. 3A,D, upper), including the bimodality in human datasets, as well as their mean dispersion relations (Fig. 3A,D, lower), can be recreated. Similarly to real data-sets, the smaller *E. coli* genome results in better coverage with similarly sized libraries (and thus higher average counts (Fig. 3B). Interestingly, in the simulated *E. coli* dataset, uncertainty is dominated by batch-to-batch variability, so detection is limited by fold-change, resulting in false negatives for genes with only small differential expression (Fig. 3C). In contrast, the lower average counts in the Human dataset mean that variability is dominated by sampling noise, and this means that false negatives are not as correlated with fold-change (Fig. 3F) and that detection is mostly dependent on mean expression rather than foldchange (Fig. 3E).

**Figure 3:**
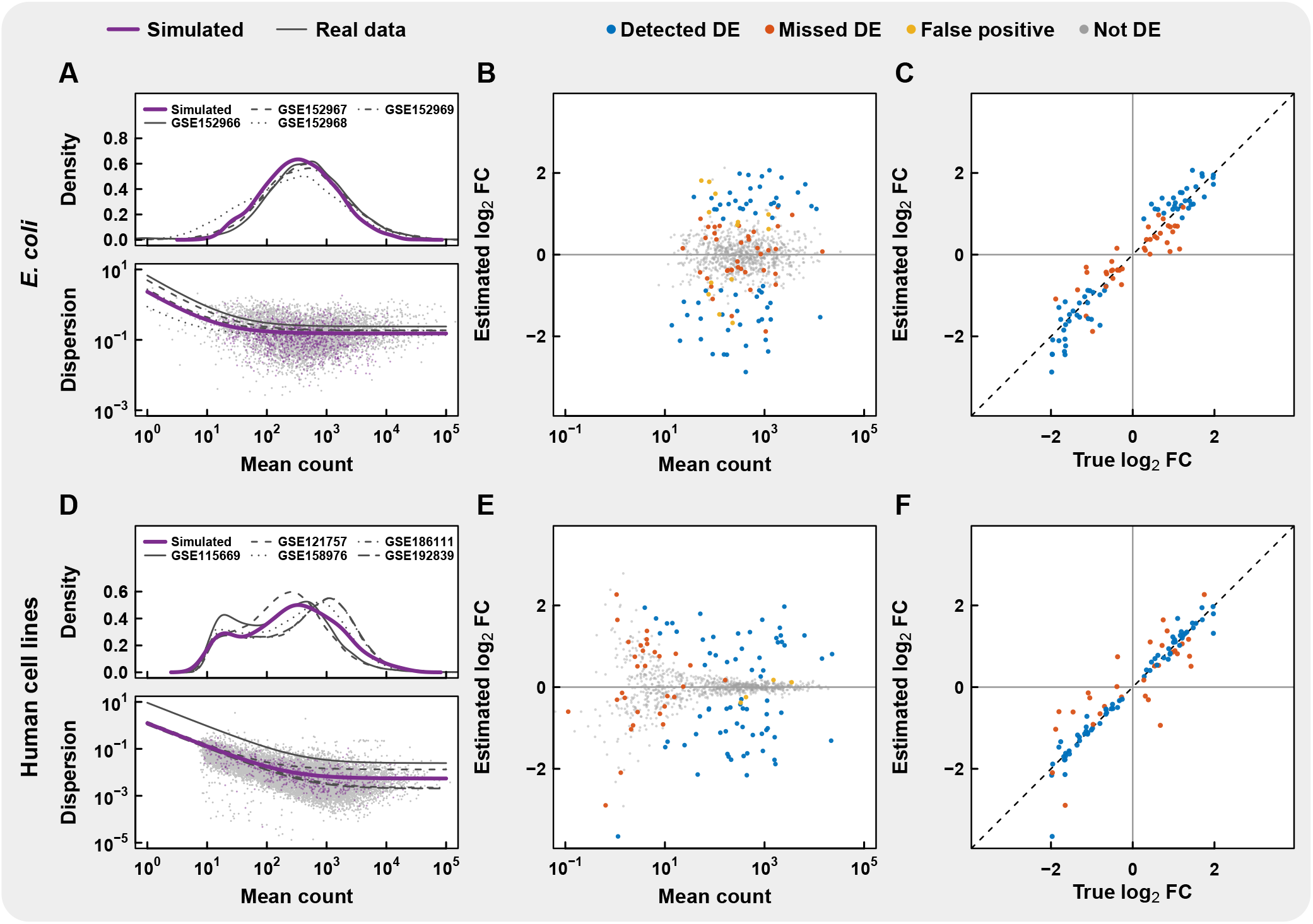
Simulated bulk RNA-seq experiments can reproduce statistics of gene expression. Two-conditions experiments were simulated with 1000 independent genes, of which 100 were set to be differentially expressed with six replicates per condition. The simulations used the *GeneRegulatorySystems.jl* prokaryote model for *E. coli* (top) and the default model for human cell lines (bottom) to generate datasets that were analysed with DESeq2. (A,D) Top, the distributions of mean count per gene are plotted along with the dispersion estimates for the simulations (purple) along with the real datasets (gray). Genes are over-dispersed compared to a Poisson model. (B,E) Minus Average (MA) plots of estimated log fold change versus the mean count, colored according to true positive, false negatives, and false positives. (C,F) the estimated log_2_ fold change plotted against the true log_2_ fold change that was set in the simulations.

### C. The model specification and scheduling system allows easy and flexible construction of simulation plans

The *GeneRegulatorySystems.jl* scheduling system allows one to organise detailed simulations of specific gene regulatory systems (Box 1). These can vary in complexity, from simply setting the initial number of molecules in the system to fully generating a simulation plan that can mimic elaborate experimental protocols with multiple instantaneous or sustained interventions, simulation branching, and sampling (with or without applying an extraction protocol).

Interventions can be used to simulate processes such as perturbation sequencing through CRISPR mutations, and can even replace the regulatory model for specific time frames. For instance, when the included templating system is used to recreate the repressilator, a classic systems biology model of genetic oscillation (*44*), simulation schedules can further be augmented with periodic interventions, such as light–dark cycles (via light-induced protein degradation).

Using the specification language, it is straight-forward to compare the undisturbed system (Fig. 4A) against explicit interventions such as express micro-injections of proteins (Fig. 4B) or their periodically induced degradation (Fig. 4C). We employ the scheduling system’s branching feature to study cell to cell heterogeneity. In particular, we investigate how multiple cells that have been synchronized through an exogenous mechanism subsequently decohere, and specifically the response of the circadian clock to exogenous signals as a form of oscillator entrainment (*45* , *46*). For example, in Figure 4 simulation branching models the divergence after collective synchronization. We quantified this process (Fig. 4D) using a short-time order parameter ((*47*) and Methods M.3). We also used simulation branching to investigate the effects of periodic interventions on the dominant frequencies of the system (Fig. 4E,F).

**Figure 4:**
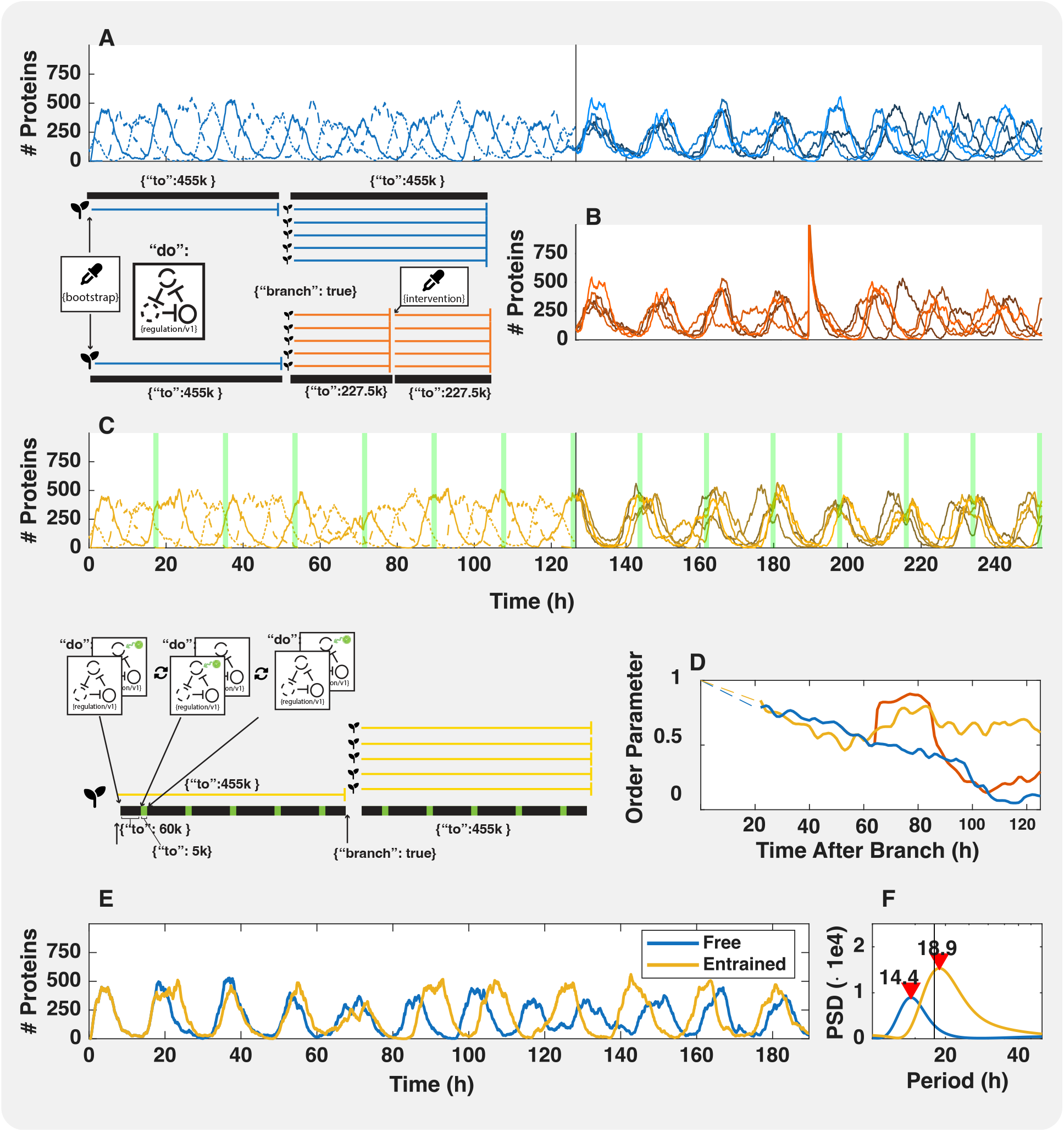
A flexible templating and scheduling system streamlines organizing simulations. The scheduling system makes it straightforward to include exogenous (potentially recurring) interventions which can be either instantaneous or sustained. In panel (**A**, blue) we simulate a repressilator which is scheduled to branch into 5 independent replicates that share their history up to the branch point (at 125 hours), after which the trajectories progressively decohere. Interventions may represent experimental actions, such as the addition or removal of reactants, or environmental influences. An example of an instantaneous intervention, the addition of a protein to the system, is shown in panel (**B**, red). Sustained interventions can also be scheduled, which can e.g. be used to mimic light induced entrainment of the repressilator by periodically swapping in a model with increased degradation of one element during illumination (**C**). Entrainment counteracts the natural tendency for signals to decohere, which can be quantified with a short time order parameter (**D**), and also slightly increases the period of oscillations from the natural frequency to the entrainment frequency (**E**). This is evidenced by a shifted peak in power spectral density from *≈*14.4 hours to *≈*18.9 hours for an entrainment period of 18 hours (**F**).

More generally, simulation branching allows arbitrary sampling of the system at various time points either from a single trajectory or from multiple independent trajectories. *GeneRegulatorySystems.jl* can thus represent both sequencing-based experimental protocols that destroy cells on measurement as well as non-destructive GFP-based cell tracing. These two cases differ in their resulting interpretation, but if the underlying dynamics are ergodic and the system has settled into equilibrium, then the two cases lead to equivalent empirical gene expression distributions. However, systems which contain feedback are not necessarily ergodic, and thus multiple trajectories have to explicitly be considered. But this comes at a considerable computational cost when attempting to faithfully represent thousands of genes in hundreds to thousands of cells. Furthermore, when modeling collections of mutually independent cells, the trajectories can desynchronize as previously shown in Fig. 4. Branching provides a convenient approach to striking a balance between single and multiple trajectory specifications, in terms of the computational trade-off as well as the synchronization issue. *GeneRegulatorySystems.jl* could in the future even be used to model nondestructive live cell sequencing (*21*).

**BOX 1**

A schedule is a type of composite model that advances the simulation through organizing multiple other models, to be invoked in sequence. Simulation results are then obtained by concatenating their individual outputs. The schedule can be specified in JavaScript object notation (JSON):

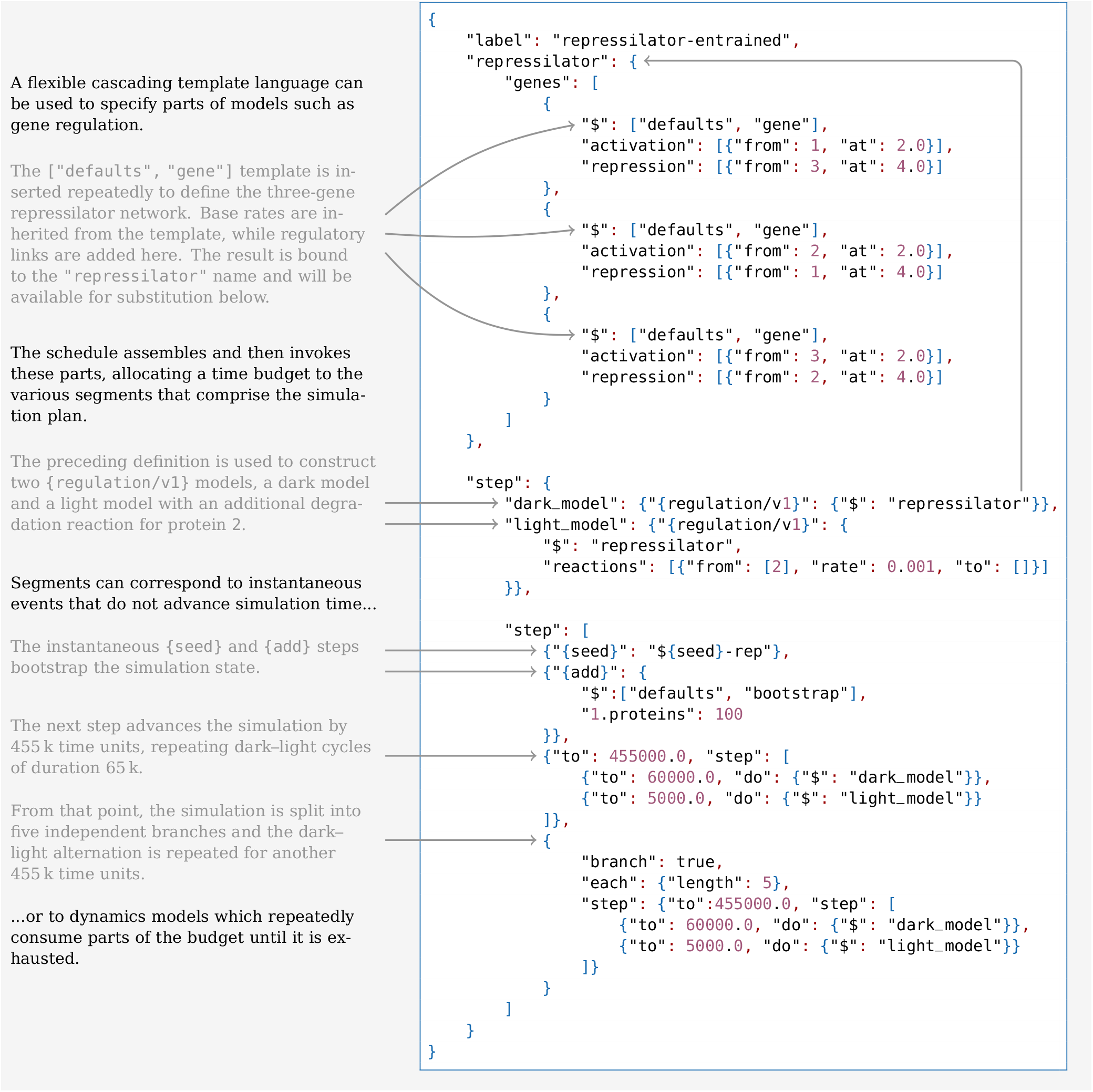

The scheduling system forms the core of *GeneRegulatorySystems.jl* as it ties the various included regulation and instant adjustment models together into reproducible experiment specifications that, besides the gene regulatory systems to be simulated, may also include initial setup and seed control, simulated omics extraction protocols, (optionally regularly repeating) interventions as well as simulation branching into independent samples sharing parts of their history.

### D. Simulation plans allow parameter and grid searches for model exploration

In dynamical systems biology applications it is often desirable to explore parametric and phase spaces in order to find boundaries between qualitatively distinct dynamical behaviors. These might include grid searches over initial conditions to find separatrices between different basins of attraction in phase space or over parameter combinations to identify bifurcations (Fig. 5). The scheduling system described above facilitates programmatic model generation so that models with different parameters can be easily instantiated.

**Figure 5:**
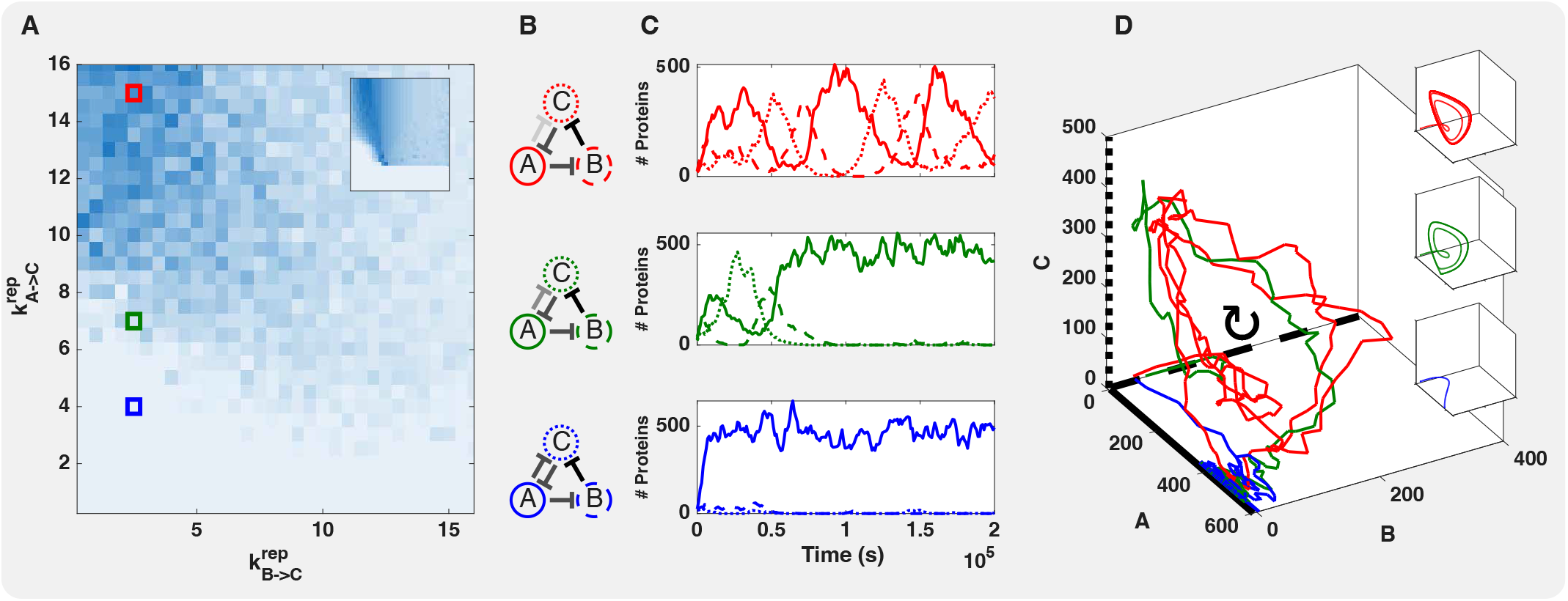
Templated plans simplify parameter space exploration and model simulation. Average (*n* = 5) peak power spectral density for a 32 *×* 32 grid search over the link strengths of repression from A to C 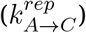) and repression from B to C 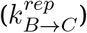), both over the range *{*0, 0.5, … , 16*}*. The inset shows the results of a similar PSD analysis on the single realization of the corresponding reaction rate equations for each parameter combination. Note that strong repression is found toward the origin. (**B**) Schematic depictions of the AC-DC models with the parameter combinations marked in red, green, and blue in (A). The shade represents the relative link strength. (**C**) Protein number as a function of time for realizations of the protein dynamics from each of the three parameter combinations showing different qualitative behaviors: oscillations (red, upper), oscillations decaying into bi-stability (green, middle), and bi-stability (blue, lower). (**D**) Phase portraits of the 3-D protein dynamics for the three trajectories shown in (C). The insets show the phase portraits for the reaction rate equations corresponding to the stochastic trajectories.

For example, in the so called AC-DC genetic circuit (*48* , *49*), which is a combination of a bistable toggle switch and the repressilator, different parameter combinations can give rise to either switch-like or oscillatory behavior. Using the peak power spectral density (Methods M.4) as a measure of oscillatory behavior, a 32-by-32 parameter grid search over half activation values for the repression from A to C and A to B was performed for 5 replicates each and averaged. This analysis reveals a wedge-like region of parameter space giving rise to oscillations (Fig. 5A). Example trajectories are shown for multiple parametrizations (Fig. 5B and also marked in Fig. 5A), corresponding to settings from the oscillatory region (Fig. 5C, upper, red), from the bistable region (Fig. 5C, lower, blue), and from near the boundary (Fig. 5C, middle, green). Fig. 5D shows the three trajectories in protein phase space.

*GeneRegulatorySystems.jl* exposes the intermediate SciML models which can then be used for further analyses. For example, models from a given schedule can be used to generate a corresponding system of continuous ordinary differential equations using the propensities to generate reaction rate equations (RREs). This functionality was used to generate trajectories as shown in the insets of Fig. 5C. These ODEs can also be used to perform grid searches for dynamical behaviors of interest (Fig. 5A, inset). It is important to note, however, that the behavior of discrete stochastic systems does not necessarily correspond exactly to their continuous RRE approximations. For example, the stochastic system in green settles to bi-stability, while the ODE system predicts persistent oscillations (Fig. 5C, inset, middle, green).

### E. Simulations can generate cell differentiation programs and cell fates

Single cell transcriptomics experiments are widely used to map the cell types that generate and constitute complex tissues into so-called cell-atlases (*50*). This approach provides a complementary approach to dissect the molecular function of organs and tissues and was used successfully to study the human brain (*51*) and lung (*52*). However, to understand in detail the processes that generate complex patterns of cell fate and differentiation throughout development, those processes need to be modeled mechanistically. Even dynamical models can be insufficient for this task if they do not represent the underlying causes faithfully (*53*), or imply oversimplified dynamics (*54*), in particular by assuming that cells equilibrate to a unique steady state, a condition that necessarily precludes the multi-stability of differentiation programs.

Since *GeneRegulatorySystems.jl* is fully compatible with feedback loops and non-linear dynamics, multiple steady states are readily realizable. As a proof of concept, we demonstrate how our software package can be used to model differentiation as an intrinsic property of the developing system itself which autonomously branches into multiple steady states using a construction inspired by the Multifate system (*55*). The multifate system is a simple differentiation program that relies on the auto-activation of genes by transcriptionally active homo-dimers, and the association of monomers into inactive heterodimers between genes. This can result in bi- or multi-stability. Utilising the reaction overlay network in *GeneRegulatorySystems.jl* allows such dimerization reactions to be specified, and the resulting circuits behave as expected, as shown by a phase portrait of the steady state for a grid search over starting concentrations of two transcription factors in a multifate-2 example (Fig. S1). In (*55*), the presence of a region of co-existence or a sharp separatrix between states can be modulated by the stability of protein dimers.

Taking advantage of this sharp segregation mechanism, we introduce a modified regulatory module in which transcription factors are proteolytically degraded by a ‘timer’ gene (with timing controlled by molecular half-life). By adjusting the decay rate of this timer, each subsequent bifurcation can be triggered at a prescribed time after the previous differentiation event (Fig. 6A, upper). Bistability can also be tuned to preferentially enter one state of the other by introducing a bias in the mutually repressive links (Fig. 6A, lower). The bistable switch along with the timer can be combined into a module (Fig. 6B), which can then be linked with two downstream modules such that the bifurcation of the upstream module triggers the start of one of its downstream timers. Thus, a simple system of such modules can be wired together to generate arbitrary trees with target differentiation timings and proportions of final cell fates, representing developmental differentiation (Fig. 6C). *GeneRegulatorySystems.jl* includes a built-in *differentiation* model template that allows such cell-fate bifurcation trees to be constructed based on user-defined timings and proportions (Methods K). The transcription factors controlling the underlying differentiation state transitions can further interact with any number of downstream genes associated with a distinct peripheral network. The template could be used to simulate realistic single cell data from complex tissue types. In many applications, synthetic data for which a ground truth is available are needed for validation. Examples include dimensionality reduction procedures, single-cellvelocity-based inference approaches (*56* –*59*), or methods used to identify transcription factors (*60*). We anticipate that this *differentiation* template will foster the construction of various types of assumed ground-truth scenarios that can be utilized for stringent benchmarking of such single-cell quantification methods. Although we have not addressed all of these use cases here, we provide an example of using simulated data to assess the sensitivity and specificity of differential gene expression calls using DESeq2 (*43*) (Fig. 3) and of clustering of cell types based on gene expression using Bonsai (*61*) (Fig. 7).

**Figure 6:**
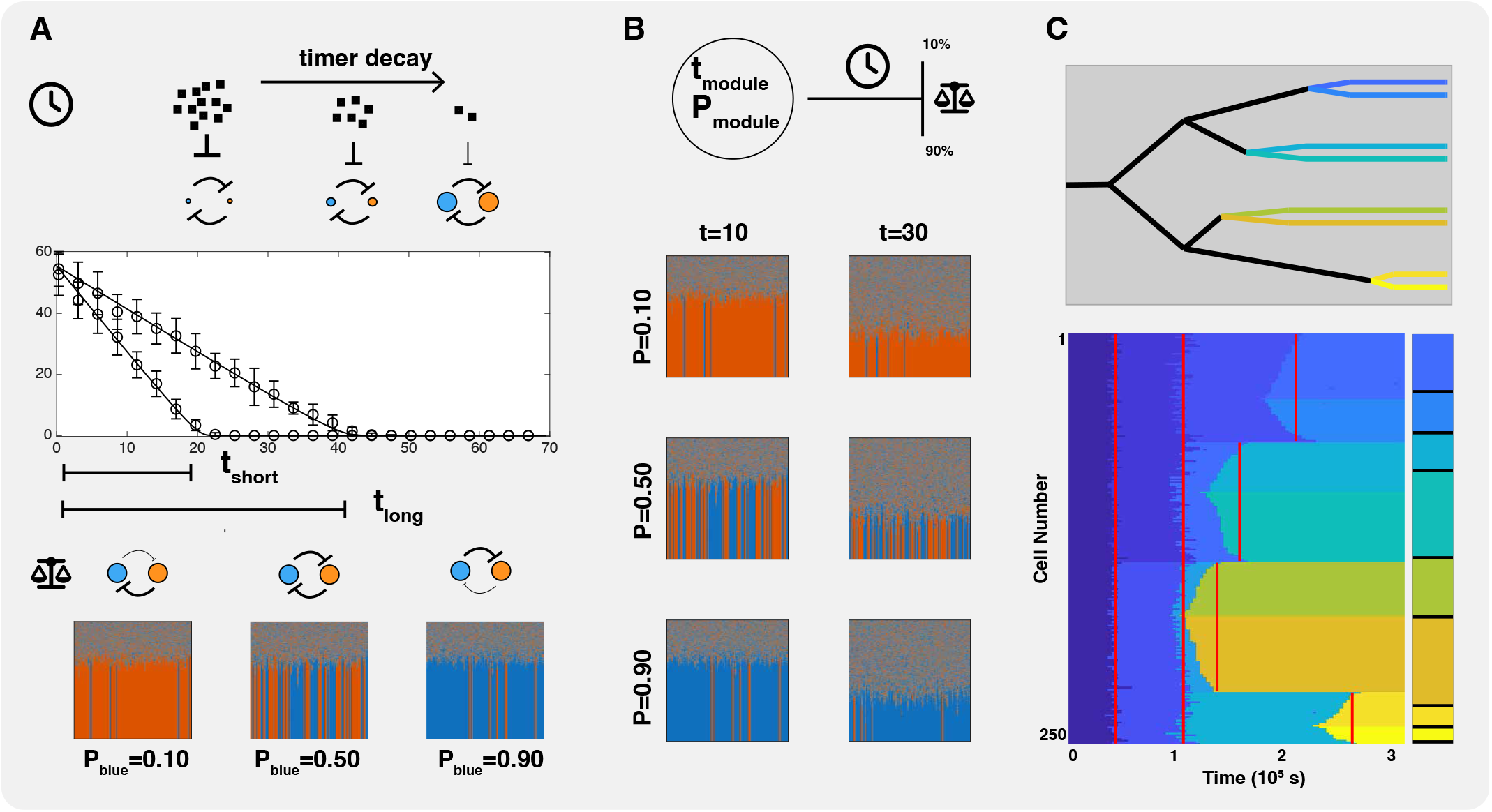
Complex cell lineages can be specified using differentiation modules. Individual modules for differentiation were designed to have programmable durations and proportions (**A**). The duration is controlled by the degradation rate of a timer protein, which reduces levels of the bistable differentiator genes’ proteins (**A**, upper). When the timer has decayed sufficiently, the system undergoes a fold-bifurcation resulting in one of two stable states. The proportion of outcomes is controlled by an asymmetry in mutual repression between the switched pair’s genes. 500 cells were programmed to give a desired proportion, and allowed to differentiate (**A**, lower). The color (orange/blue) represents the enrichment of that differentiator gene relative to its sibling. Modules can be programmed to generate a specified duration and proportion of states (**B**). These modules can then be combined to design complex lineage trees (**C**, upper). 500 identical cells were instantiated, and allowed to differentiate with the designed program (Methods K). The color in (**C**, lower) indicates which combination of differentiator genes are active in each cell. The bar to the right indicates the target proportions for each cell type.

**Figure 7:**
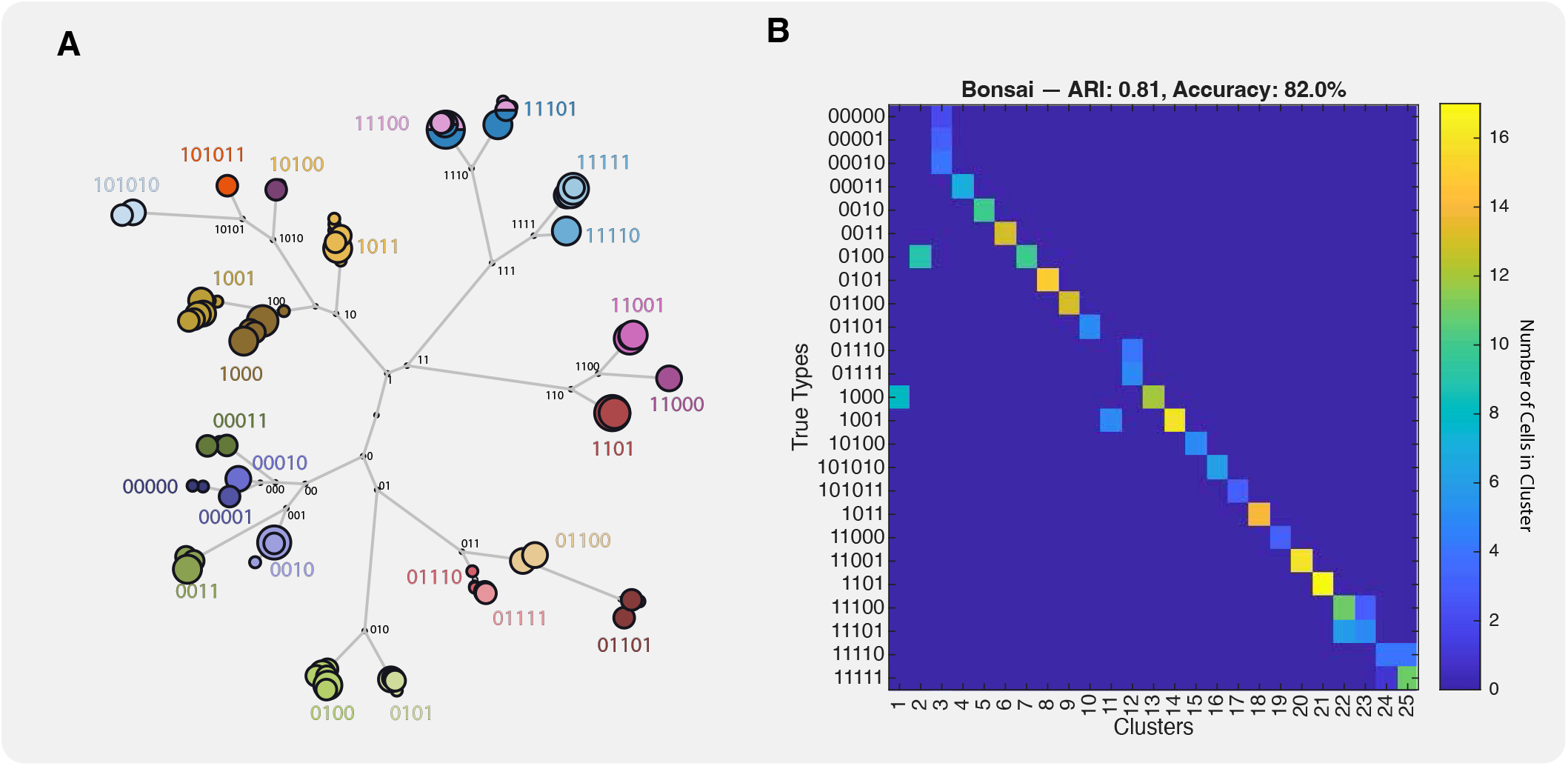
Simulated differentiation can be captured by state-of-the-art third-party inference tools such as *Bonsai*. A schedule utilizing the random differentiation template (Methods L.2) with 25 terminal states was simulated (**A**). The final cell state can be indexed by inspecting which transcription factor in each bistable pair is active. Bonsai was used to infer a tree structure from mRNA counts, and plotted using Bonsai-scout. Nodes were colored by their true identity. Confusion matrix comparing clustering assignments (columns) with ground truth cell types (rows). Color intensity indicates cell count. Statistics are the Adjusted Rand Index (ARI) which quantifies whether pairs of cells of the same type in the ground truth are in the same cluster in the analysis, and the classification accuracy, which is the fraction of correctly identified cells based on an optimal assignment of clusters to true types.

### F. Simulations can generate data for downstream applications

We can explore the space of differentiating systems according to this setup by randomly sampling the differentiation tree, peripheral network and interlink structure, and then studying the trajectories generated from such realizations. *GeneRegulatorySystems.jl* provides a *random differentiation* model template (Methods L.2) implementing this idea for a specific choice of structural parameters for these components. This allows for the convenient construction of large regulatory systems with thousands of genes that intrinsically show differentiating behavior.

Parameterizing the template with a random seed, the construction becomes fully deterministic and can be instantiated into experiment schedules like any other model. During schedule execution, the high-level stochastic template is automatically lowered in a multi-stage procedure down to the jump process definition from which trajectories can be directly sampled as described above. Due to the deterministic nature of this construction, the intermediate representations (which describe various structural aspects of the regulatory system such as the differentiation tree of the backbone), need not be retained explicitly because *GeneRegulatorySystems.jl* can fully reify them from only the schedule specification and the random seed, both of which are automatically included in the simulation outputs. The resulting sampled trajectories can then be used for downstream analysis.

In this way, *GeneRegulatorySystems.jl* can be used to generate synthetic gold-standard singlecell and bulk multi-omics data with completely known intrinsic structure, which may, for example, be used to validate methods for technical noise reduction, RNA-velocity, pseudotime and GRN inference. We include canonical experiment schedules that define differentiating systems of various sizes with the software package. As an example, we tested this with Bonsai (*61*), a new state of the art method for inferring lineage trees which relate any high-dimensional objects, such as single cell transciptomes (Methods M.5). For this purpose, we created a synthetic dataset using the random differentiation model for 25 target states with 256 peripheral genes which are regulated by, but have no regulatory feedback to the central differentiation tree. Synthetic mRNA data was pre-processed with Sanity (*62*) and a tree was inferred with Bonsai (*61*) (Fig. 7A). Informed by this tree, Bonsai was then used to cluster the genes. For comparison, uniform manifold approximation and projection (UMAP) (*63*) was applied to the same dataset, and k-means clusters were then obtained. Bonsai performed better (Fig. 7B) than UMAP in both cases, and having the full set of genes available improved correct cluster identification, but only very slightly. UMAP clustering was not as good, and interestingly, UMAP performed better on peripheral genes alone than on the entire dataset.

**Table 1:**
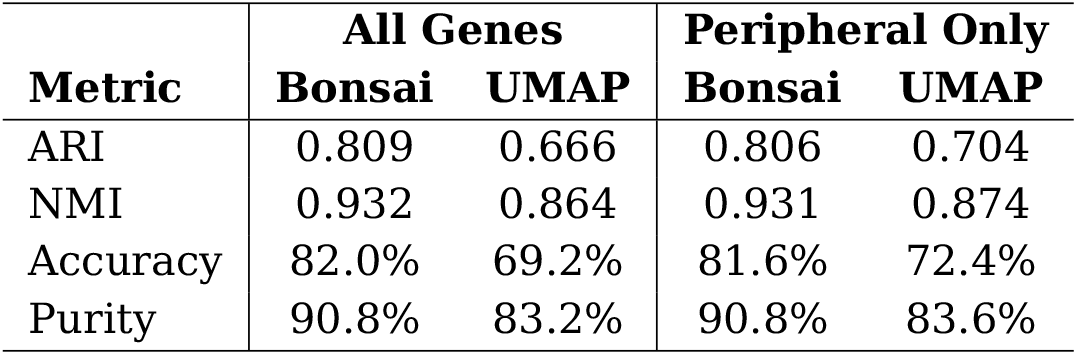
Side-by-side comparison of clustering performance metrics for Bonsai and UMAP with K-means clustering (*k* = 25) for all genes (*n* = 304) or only the peripheral genes (*n* = 256). See Methods M.5.

|  | **All Genes** |  | **Peripheral Only** |  |
|---|---|---|---|---|
| **Metric** | **Bonsai** | **UMAP** | **Bonsai** | **UMAP** |
|---|---|---|---|---|
| ARI | 0.809 | 0.666 | 0.806 | 0.704 |
| NMI | 0.932 | 0.864 | 0.931 | 0.874 |
| Accuracy | 82.0% | 69.2% | 81.6% | 72.4% |
| Purity | 90.8% | 83.2% | 90.8% | 83.6% |

### G. Exact simulations can be scaled to tens of thousands of genes

We demonstrate in principle that *GeneRegulatorySystems.jl* allows for the simulation of genome-scale regulatory networks with eukaryotic kinetic rates, without compromising on mechanistic detail. We simulated networks of up to 32,768 genes from the Kronecker template (Methods L.1) for 172,800 s (two days) of model time (Fig. 8). All benchmarks were run single-threaded on a virtual machine (Intel Xeon Gold 6230R CPU, 2.1 GHz base frequency, no swap, running Ubuntu 24.04). For the largest tested network, the exact RSSA-CR algorithm (*64*) took 2239s to run one trajectory, compared to 566s when using the tauleaping approximate method (*65*). We compared *GeneRegulatorySystems.jl* performance against COPASI (Python based, (*66*)) and dyngen (R based, (*67*)), exporting the generated reaction system in the SBML format (*68*) for those other tools to use the same chemical reaction network. Ours is the only tool of the three that is able to scale exact simulations beyond thousands of genes, but dyngen is competitive when utilizing tau-leaping, running the largest network in 762s (Fig. 8A). Our tool incurs a high base cost for model build time (50 s) and peak memory usage (2 GiB), mainly due to the Julia JIT compilation, but this overhead is paid only once per session, and our tool has better scaling for large networks (Fig. 8B and C). For a fair comparison, we also benchmarked using dyngen’s 10,000 gene regulation model as a reference (which uses a quasi-steady state promoter representation), exporting it to our engine, in the regime of which their tau-leaping method is faster than ours (26.7s vs 40.7s), but our best exact SSA method remains tractable (884.4s) (Table S2).

**Figure 8:**
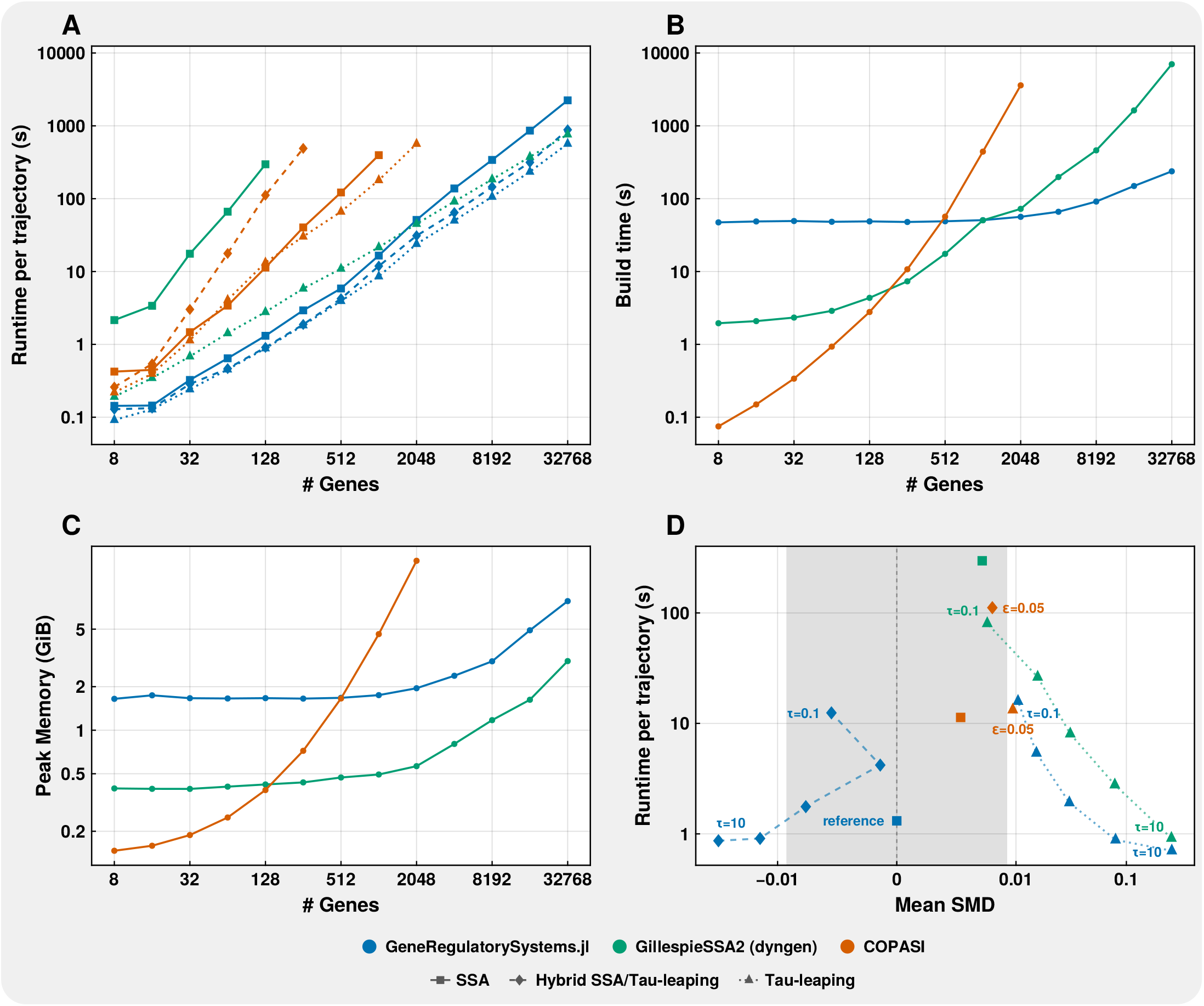
Performance comparisons across different simulation engines demonstrate the advantages of *GeneRegulatorySystems.jl*. Networks of 8 up to 32,768 genes, generated using the Kronecker template (Methods L.1), were exported to other simulation frameworks such that the concrete chemical reaction networks were identical. The largest network contains 196,611 molecular species and 327,680 reactions, 65,536 of which have state-dependent rate coefficients. (**A**) Simulation runtime, averaged over three trajectories, each spanning 172,800 s of model time, for exact SSA, tauleaping (*τ* = 3 for dyngen and *GeneRegulatorySystems.jl*; COPASI adaptive with *ϵ* = 0.05), and hybrid methods that fall back to exact SSA for critical (low copy number) reactions (App. A). (**B**) Time to construct the reaction networks and prepare the simulation. (**C**) Peak resident set size (RSS) during build and simulation. (**D**) Runtime against bias (signed SMD, Methods M.6) for the approximate methods on the 128 gene network, against an exact RSSA-CR reference (*n* = 200). The gray band is the inherent sampling noise (95th percentile of |SMD| across reference splits). Fixed-step methods use *τ ∈ {*10, 3, 1, 0.3, 0.1*}*. COPASI’s tau-leap methods use *ϵ* = 0.05.

Tau-leaping is an approximate method and so it generally fails to reproduce the CME distribution, especially for low-copy-number species, but there are variants which can mitigate this (as detailed in App. A). For models with explicitly represented binary promoter states, we found that running critical tau-leaping (*69*) (referred to as hybrid SSA/tau-leaping) with control parameter *n*_*c*_ = 2, so that the promoter switching reactions are simulated exactly while the other reactions are tau-leaped, has a competitive runtime (882s for the largest network), while in principle maintaining high fidelity. We illustrate this tradeoff in Fig. 8D, for the 128 gene network, where it remains tractable to run many exact simulations and quantify method fidelity empirically. COPASI’s tau-leaping is adaptive only, and its hybrid SSA/tau-leaping method hard-codes *n*_*c*_ = 20. dyngen supports only fixed-step tau-leaping with negative counts clamped to zero.

To show the necessity of the explicit promoter switching model, we measured the fidelity of the faster quasi-steady state (QSS) approximation for the 512 gene network for different switching rates (Fig. S3). We found that “bursty” regimes with slow switching (characteristic of eukaryotic systems) are poorly approximated by the QSS model (0.48 histogram distance against 0.27 noise floor; Methods M.6), while running at comparable speed, corroborating previous studies (*70*). On the other hand, fast promoter switching is expensive to simulate explicitly: promoter switching events raise the total from 4.3 *×* 10^7^ to 1.6 *×* 10^8^ events per trajectory, while the QSS approximation is indistinguishable from the telegraph model in this setting. This comparison underscores the need for a flexible specification system, allowing for model reduction when appropriate.

## Discussion

The current understanding of how gene regulation organizes cellular function is based on both detailed molecular mechanisms of single-gene regulation and statistically descriptive models of large-scale genomic patterns. Associational descriptions such as gene co-expression clustering are often used for predicting cell function and have proven useful when employed to identify possible targets for more focused experiments. However, associational descriptions alone lack a causal interpretation and thus do not faithfully capture the behavior of biological systems under intervention, in general, unless they are mechanistically modeled.

A major remaining challenge in complex systems biology is how to compile a molecular-scale causal understanding of gene expression regulation into systematic dynamical models at the genomic scale. *GeneRegulatorySystems.jl* is our proposal to bridge this divide by assembling the detailed knowledge resulting from decades of prior research on the biophysics of gene regulation into large-scale cell-level simulations. Our simulations can capture both dynamics of and interventions on gene regulation to mirror realistic experimental designs and generate “gold standard” *in silico* data that can be related to experimental omics data. We envision that *GeneRegulatorySystems.jl* can be the basis for causal predictions of global network effects of local interventions, which can lay the ground-work for general causal inference in biological systems.

Inferring a GRN from expression data neces-sarily implies a dynamics model, even if one is not explicitly stated. Even simple correlation-based GRN inference implies dynamics where each gene’s expression is only a linear combination of the other genes, and the transcriptome therefore reaches a unique steady state. Often, the dynamics associated with an inference method are left implicit or vague, and this precludes their use for causal, mechanistic reasoning. It is always possible to infer *a* GRN from expression data, but the flexibility afforded by under-specified dynamics generally means that its model structure does not agree with that of the biological ground truth. In short, this is a case of model over- or under-fitting, which is typically mitigated by appropriately constraining a model. For GRN inference, that would mean constraints which accurately reflect true biological mechanisms.

Our best mechanistic understanding of gene regulation comes from quantitative characterizations of the expression of single genes. We formulate this detailed knowledge into singlegene reaction cascades that accurately reproduce well-known behaviors at this level, such as expression burstiness and expression mean– variance relationships. We then assemble these components into a cell-scale chemical reaction network whose dynamics interpretation is given by an associated chemical master equation; this is a generative model of whole genome expression dynamics which we refer to as a gene regulatory system rather than a gene regulatory network because the dynamics aspect is essential for understanding it.

We are not aware of simulators of gene regulation and expression with gene architectures as realistic as the one used in *GeneRegulatorySystems.jl* that can also scale to tens of thousands of genes. Previous efforts to apply such models at the genomic scale either abandon exact stochastic simulation altogether for approximate techniques such as a reaction rate formulation (ODE or CLE) (*17* , *54* , *71* , *72*) or tau-leaping (*65* , *67*), or they apply exact SSA methods to unrealistically simplified models that are no longer mechanistically faithful, for example, models which do not reflect the discrete nature of promoter state switching (*67*). All of these simplifications can result in quantitative and qualitative differences from the dynamical behavior given by a discrete-state master equation, especially in the regime of small numbers of molecules, which is often true of transcription factors with single-digit counts per cell (*73*).

Conversely, tools that can represent comparable mechanistic detail target smaller circuits. For instance, GRN_modeler (*74*) provides an expressive toolkit for assembling networks from customizable gene templates, and also supports spatial modeling, but it uses COPASI (*66*) internally, which does not scale to large systems, as our benchmarks show.

It is now possible to scale the exact formulation directly to larger systems, in part due to the off-the-shelf availability of more efficient simulation algorithms and also faster hardware, made accessible by convenient integrations such as ModelingToolkit.jl (*75*) and the entire SciML ecosystem. While these tools greatly facilitate turning a model into a simulation, the model itself still must be assembled for a large network of co-regulating genes where each gene has the realistic architecture described above. *GeneRegulatorySystems.jl* facilitates the assembly of cell-scale networks from single-gene expression cascades, and in this way extends dynamic biophysics models from genes to genomes.

However, the behavior of a gene regulatory system, or even a single gene, is not context free; it depends crucially on the details of the experimental design. We provide a flexible scheduling system by introducing a grammar of *in silico* experiment description that abstracts and generalizes the bespoke designs previously hard-coded in software such as dyngen and others (*67* , *76*). As an analogy, if an experimental design is a sentence, previous tools defined a sentence structure and allowed the user to replace only certain words, while our experimental grammar allows much more flexibility in sentence construction, provided each sentence is syntactically correct. When invoked, the scheduler then interprets this experiment specification and organizes its atoms, the simulation segments, into complete and reproducible simulations yielding structured and standardized datasets. This ability to generically formulate diverse experimental settings is supported by a convenient templating scheme. Collectively, the scheduling system allows the user to include interventions, grid searches, and modeling of technical and biological replicates, which not only allows simulations that are much more closely aligned to real experiments and data, but also facilitates simulating novel conditions for which experimental data do not yet exist. *GeneRegulatorySystems.jl* and its accompanying graphical user interface (GUI) thus provide a programming language and environment of *in silico* experimental design that can capture gene expression dynamics for entire genomes. In this study, we highlight the versatility of this design language by producing rich temporal datasets for classic wet-lab experiments on simple systems known from molecular biophysics, developmental genetics, and synthetic biology. We also assemble these smaller components into larger systems, such as those that exhibit robust cellular developmental differentiation. In combination, this means that *GeneRegulatorySystems.jl* employs innate principles of developmental organization to channel the dynamics of non-linear systems to form cell fate maps and thus achieve emergent behaviors analogous to patterns found in nature. In conclusion, our results demonstrate the power of the framework for inspiring wet-lab design of diverse gene regulatory systems and their study through realistically simulated experiments. We envision that our simulation approach can enable the inference of parameters of structural models from data in a way that allows subsequent design of new experiments through an iterative feedback-cycle.

## Methods

### H. A programming language for simulating gene regulation

In *GeneRegulatorySystems.jl*, simulations are orchestrated by a flexible scheduling system. Its primitives are models that define how to advance the simulation state, either instantaneously or progressing along the simulated time dimension. Each dynamic regulation model is user-specified by a skeleton GRN detailed by specific reaction base rates, and optionally extended by additional mass-action reactions. The package also includes sensible base rate defaults and model templates to automatically assemble larger regulatory systems from more abstract specifications. Specific regulation models are organized into schedules, which control the simulation by ordering, compiling, and executing their contained models, recording results, and reporting on progress. Since the default mode of operation is sampling from stochastic models, this also includes careful control of the PRNG to ensure the computational reproducibility of all simulations.

When models are invoked to progress the simulation, they are given a limited time budget restricting how far along the simulation time dimension they may advance. This also applies to schedules, which are themselves implemented as composite models that keep invoking their elements in sequence, forwarding the remaining budget until it is exhausted. Thus, schedules can be elements of other schedules.

All models are assembled according to userdefined specifications and are composed of building blocks from a pre-defined (but extensible) library. For schedules, these specifications are nested and will be interpreted recursively when invoked. The scheduling subsystem thus provides a domain-specific programming language with its semantics defined in terms of the compilation and budget-restricted invocation of the orchestrated primitive models, as individual simulation segments. Specifications can be encoded in JSON, which acts as the schedules’ source code. The scheduling language is deliberately Turing-incomplete, but it does support the repetition of schedules as long as each step consumes a fixed part of the (finite) time budget, or alternatively the iteration of a (possibly implicitly) specified finite list of steps. This construction guarantees that the order and execution context of all simulation segments is statically determined and each segment can be addressed by path and reified individually for analysis.

Subject to flexible output control, parts of the realized trajectories are collected in long-style Apache Arrow (*77*) format and persisted along with all originating specifications, randomness seeds, and a manifest of the software environment to support the full reproducibility of the results, even of individual segments in isolation. For user convenience and to avoid excessive redundancy in the specification of complex schedules, the language includes a templating facility for lexically scoped reference substitution. The language further supports a branching mechanism that allows the realization of independent trajectory copies from a specific time point onward, optionally with variant models. In combination, these features allow for the specification of various experimental setups, including grid searches over a parameter space.

### I. Graphical user interface

*GeneRegulatorySystems.jl* features a graphical interface with a schedule editor and a reactive and interactive visualization of the network model and the simulated output, so that the user receives instant feedback when making changes to the specification. This responsiveness facilitates rapid research iteration, enabling exploration of how model structure and parameter choice translates into different dynamical outcomes. The interface is implemented as a Pluto.jl notebook (*78*), which makes it easy to tailor the application for specific modeling purposes and extend the dashboard with downstream analyses. Being web-based, it also allows for decoupling the interface from the simulation engine, so that the latter could run on high-performance computing clusters.

The application uses Cytoscape.js (*79*), a WebGL-based rendering engine, enabling the visualization of large-scale hierarchical networks of thousands of genes, as well as the chemical reaction network underlying the regulatory networks. We have also optimized the rendering of the simulation output to allow the inspection of trajectories with many events and branches.

### J. A baseline model template of intra-cell gene regulation

All simulations so far are based on a regulation model template that allows instantiating distinct genes and connecting them by a regulatory network. Each gene is modeled by a reaction cascade that starts with a binary state-switching abstracted promoter region which, when it is in its active state, forms elongation complexes to be successively transformed by transcription, processing (such as splicing) and translation into proteins. The intermediate products may decay spontaneously. These reactions are mediated by shared resources, specifically RNA-polymerases, ribosomes and proteasomes.

Transcriptional regulation between genes is possible by any species (typically proteins) acting as transcription factors that temper the promoter switching rates, respectively for activation and repression. This assumes a separation of time scale between TF binding and the resulting changes in effective promoter configuration. If a single gene is affected by multiple regulators of the same link type, those influences are aggregated by a generalized mean, which interpolates smoothly between their minimum (the default) and maximum and can also be set to their geometric mean.

The model can also be extended by an overlay network of additional mass-action reactions to express further regulatory mechanisms. In addition, the template includes a post-transcriptional repression mechanism via targeted proteolysis, implemented as purely mass-action reactions.

A network with *n* genes contains at least 5*n* molecular species, plus up to 3 shared resources, and potentially more if specified in any overlay reactions or if any genes’ inactive promoters are included in the representation. The molecular species are linked by 8*n* + *k* (“purely”) mass-action reactions with constant rate coefficients, where *k* is the number of specified proteolytically repressive links and the total number can again be increased by overlay reactions if specified, and further 2*n* reactions with statedependent rate coefficients aggregating the inbound transcriptional regulation for each gene (details can be found in App. B.).

Default reaction rate constants are set based on (*80* , *81*), but may be freely redefined. All parameters are configurable by gene or link. When the model template is instantiated with concrete parameters, including the GRN, it is automatically compiled to a Catalyst.jl ReactionSystem, imbued with a pure-jump-process CME interpretation and then further processed to obtain a concrete JumpProcesses.jl JumpProblem to be integrated.

By default, trajectories are sampled exactly via RSSA-CR (*64*), or a sorting direct (Gillespie) method for smaller systems (*82*). Any nonzero initial states can be provided through the scheduling mechanism as instantaneous adjustments, i.e., as user-defined interventions at time zero. Recording of results can be regularly spaced, continuous, or a combination thereof. At scheduled time points (typically at the end of trajectories), the recorded state can optionally be processed by a simulated extraction scheme, either assembled from provided subsampling building blocks or from predefined extraction schedules. The latter include an idealized PCR protocol where the results are filtered for transcripts, then sampled without replacement, accumulated through multiple rounds of i.i.d. duplications or removals without replacement, and finally sampled once more without replacement to achieve a predefined target total count.

### K. Differentiation model template

We use the baseline model template to assemble more complex regulation models. In particular, we provide a higher-level *differentiation template* that channels trajectories into distinct cell states according to a user-defined differentiation tree with specified branch times and split ratios. The construction of each of its inner nodes is based on a switching motif of two mutually repressive *differentiator* genes, which further are initially fully repressed by an upstream gene. When the cell has acquired the corresponding upstream intermediate state, that latter gene is itself repressed and its product starts decaying, thereby acting as a molecular timer. Once the timer is sufficiently depleted, the switch will collapse into one of its two modes. To approximately achieve the target onset delay and outcome ratio specified by the user, the genes’ parameters are adjusted using coefficients determined from a previous calibration experiment, which can be found in the *GeneRegulatorySystems.jl* repository (cf. also Fig. 6). The differentiation tree can optionally be configured with an additional peripheral GRN, and arbitrary inter-regulation can be specified.

### L. Random model templates

We further provide GRN templates from which specific GRSs can be sampled at random, distributed according to an imposed network model. The other required parameters are then specified as userconfigurable probability distributions, or alternatively as fixed values, in the concrete simulation schedule. For all presented simulation experiments, we configured the genes’ base rate parameters to use fixed values.

All templates are instantiated with concrete parameters, which are always specified as part of the enveloping schedules, including any randomness seeds. Given any schedule and the specific *GeneRegulatorySystems.jl* version, the resulting networks are therefore reproducible at all levels of abstraction, and available for analysis using the reification tooling.

#### L.1 Kronecker template

A simple variant implements a stochastic Kronecker graph (“SKG”)-based linkage (*83* , *84*) independently per regulatory link type, with the genes’ and links’ parameters chosen i.i.d. according to configurable univariate distributions; this setup is available as the *Kronecker template*. The SKG construction is based on a small “initiator” matrix of probabilities, which is then expanded by taking its Kronecker power to obtain a larger random matrix. Sampling i.i.d. from the latter’s entries produces the network’s adjacency matrix. If requested in the model specification, the initiator is optionally adjusted by adding a common offset to its entries on the logit scale. The offset value is found by an optimization procedure to achieve a target inbound link count per gene in expectation.

SKGs have the small-world property (*85*), and a nested community structure which is self-similar at multiple scales (*83*). A procedure for fitting the initiator to a given network is available (*84*) so that SKGs could be used as a structural prior in inference applications. However, they should not be expected to reflect the presence of feedback motifs accurately. Further, SKGs are not scale-free for all parametrizations (*86* , *87*) as their node degree distributions can deviate from the power-law shape due to repeated factors in the expansion. In this work, SKGs are rather used to produce larger GRNs from a small number of structural parameters for benchmarking purposes as a more flexible alternative to the well-known preferential attachment model (*88*). SKG sampling scales well to larger graphs (*89*).

#### L.2 Random differentiation template

These ideas are combined in the *random differentiation template*, which assembles a core and a peripheral network, respectively based on the differentiation and Kronecker templates, and then connects them through an *interlink template*. The interlink is here restricted to transcriptional regulation directed from core to periphery. Each peripheral gene has the number of inbound links for both link types sampled from configurable distributions, the corresponding links’ sources are chosen i.i.d. from the core network’s differentiator genes, and their parameters i.i.d. from configurable distributions as well.

The target timings and other parameters of the differentiation’s tree nodes are chosen i.i.d. from configurable distributions, while the target split ratios are determined indirectly by the following scheme: First, the terminal states’ target proportions are sampled from a (configurable) multivariate distribution and potentially normalized to obtain a probability vector. This vector is then agglomerated into a differentiation tree by the well-known Huffman algorithm to achieve a minimal expected count of intermediate cell states.

## M. Analytical Methods

### M.1 Fano factor

The Fano factor quantifies the deviation of mRNA expression variability from Poissonian behavior and was calculated as the ratio of variance to mean

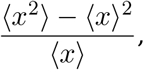

where x represents the counts of mRNA molecules. Values equal to 1 indicate Poisson statistics (typical of constitutive expression), while values greater than 1 indicate the super-Poissonian noise characteristic of bursty transcription.

### M.2 Bulk RNA-seq simulations

Each gene in the simulation (cf. Fig. 3) is an independent, unregulated gene expression cascade for which the translation rate is set to 0. For the prokaryotic model the average RNA lifetime was set to be 100 s and for the human simulation the average processing time was set to be about 17 minutes and the mRNA lifetime to about 8 hours. Ten percent of the genes were chosen to be differentially expressed with a log_2_ fold change chosen from a uniform distribution between 0.25 and 2 (1.19x to 4x) tuned by adjusting the transcription initiation rate. Each sample consists of 50 independently simulated cells and data was collected after a burn in time of 10 mRNA lifetimes using the simulated PCR extraction model. In real RNA-seq data most variation comes from batch effects. To simulate this, the base transcription initiation rates for all genes in a sample were scaled by an independent log-normal factor with mean 1. The dispersions for the simulated data were chosen to replicate real data by adjusting burst statistics using the promoters’ activation and de-activation rates. For comparision, the following datasets were chosen.

- E. coli: four clinical isolates grown in either RPMI or in human serum, 6 vs 6 replicates (GEO GSE152966–GSE152969).
- Human: five cell-line perturbation experiments (GSE115669, GSE121757, GSE158976, GSE186111 and GSE192839)

Simulated and real data were both analysed with DESeq2 1.46 (R 4.5.1), using the default settings. Significance was determined using a Benjamini–Hochberg adjusted p-value set at *P <* 0.1 (default threshold value for this procedure). For the dispersion comparisons, human genes were pre-filtered for *≥* 10 counts.

### M.3 Order parameter

The order parameter quantifies the degree of synchronization or coordination among multiple temporal gene expression trajectories. It measures the ratio of between-trajectory variance to within-trajectory variance, ranging from 0 (independent, unsynchronized dynamics) to 1 (perfectly synchronized dynamics). For a set of parallel trajectories *x*_*i*_(*t*), the order parameter was calculated using a sliding window approach:

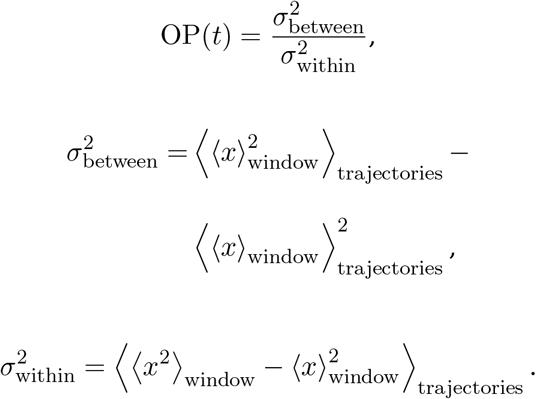

In this work, we select a window size of 20 hours.

### M.4 Power spectrum analysis

Power spectral density (PSD) analysis was performed to characterize the oscillatory dynamics of protein expression in the repressilator system. The PSD quantifies the distribution of signal power across different frequencies, revealing dominant periods in the time series data. Using MATLAB, the signal was interpolated onto a regular time grid using interp1 (linear interpolation), with any remaining NaN values set to zero. The interpolated signal was mean-centered. The PSD was computed using pspectrum with frequency limits of [0, 3 *×* 10^*−*5^] Hz and frequency resolution of 6 *×* 10^*−*6^ Hz. Dominant oscillation peaks were identified using findpeaks with a minimum peak height threshold of 1000 and constrained to frequencies *f <* 2 *×* 10^*−*5^ Hz.

### M.5 Clustering analysis

To evaluate the performance of the clustering algorithms, goldstandard expression counts were generated from a system that autonomously differentiates into one of 25 terminal states. Connected to this network were 256 peripheral genes which had no regulatory feedback into the core differentiation network. These dynamics were realized in 250 independent cells, and each cell acquired one of the 25 possible states. This state was recorded as the ground truth. The RNA expression data at the final time point was collected through simulated PCR extraction and analyzed either using Bonsai or by UMAP. For Bonsai, the counts were pre-processed using Sanity and then run with Bonsai’s default settings. The clustering was done using the clustering functionality in BonsaiScout. Similarly, UMAPs were constructed using the python *umap* package with default parameters and k-means clustering was done using the scikit learn clustering package. Both clusterings were done with *k* = 25 target clusters. For each clustering the following statistics were calculated.

#### Adjusted Rand index (ARI)

Considers every pair of cells and checks whether both partitions (clustering and ground truth) agree on grouping them together or apart. Adjusted for chance so that a random clustering scores 0. Range: [*−*1, 1], where 1 means perfect agreement (*90* , *91*).

#### Normalized mutual information (NMI)

Measures how much knowing the cluster assignment reduces uncertainty about the true type (and vice versa), normalized by the entropies of both partitions. Range: [0, 1], where 1 means that the two partitions carry identical information (*92*).

#### Accuracy

The fraction of cells correctly classified after finding the optimal one-to-one mapping from clusters to true types using the Hungarian algorithm (*93*).

#### Purity

For each cluster, the fraction belonging to its most common true type, averaged across all cells. A clustering with many tiny pure clusters scores high on purity but low on ARI (*94*).

#### M.6. Accuracy measurement

We quantify the error of approximate methods against the exact SSA reference via the following distance metrics over the joint probability distribution 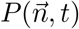 of the CME, where 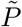 is the distribution sampled by the approximate method, 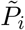 and *P*_*i*_ are the marginal distributions of species *i, T* are the 10 sampled time points, and *σ*_*P,i*_(*t*) is the reference standard deviation. The sampling noise floor is estimated as the self-distance of bootstrapped samples of the reference.

#### Histogram distance: (*95*)

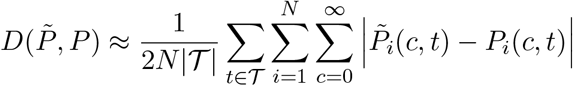

#### Signed standardized mean difference (SMD)

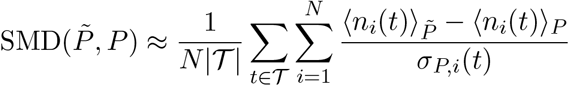

## Acknowledgments

We thank D. Weigel for his generous sponsorship, discussion, and overall support of this work. D.J. thanks E.A. Miska for his support. We thank K. Muandet, M. Munkhoeva, T. Narendra, G. Schweikert, I. Moutsopoulos, I. Mohorianu, N. Kalábová for discussions at the early stages of this project. The authors also thank the members of the Drost, Weigel, and Miska laboratories for fruitful discussions and feedback.

## Authors contributions

H.-G.D. conceived and led this study. D.J. and L.M. derived the models underlying *GeneRegulatorySystems.jl*. L.M. implemented *GeneRegulatorySystems.jl* with feedback from D.J., S.M., H.-G.D. S.M. implemented the graphical user interface (‘GUI’) for *GeneRegulatorySystems.jl*. D.J. performed simulations using *GeneRegulatorySystems.jl* and assembled the figures with feedback from all authors. Analyses were performed by D.J. and L.M. Benchmarks were performed by S.M., adapting earlier work by L.M. All authors interpreted and discussed the results. D.J., H.-G.D., and L.M. wrote the manuscript with contributions from S.M. All authors have read and approved the final version of the paper.

## Data, code, and materials availability

Our simulator *GeneRegulatorySystems.jl* is available as Open Source Software via GitHub at https://github.com/drostlab/GeneRegulatorySystems.jl. Scripts and simulation schedules to reproduce the experiments and results presented in this study can be found at https://github.com/drostlab/grs-paper, along with the Julia and *GeneRegulatorySystems.jl* versions used to produce the results.

## Supplementary Notes

### A. The chemical master equation

The molecular state of a cell can be represented by the number densities of the molecules that comprise it at a given time *t*. While the state of the cell, in principle, consists of all of its molecules and metabolite constituents, in practice, it is often macromolecules, the RNA transcripts and the proteins, that are of interest for investigating the evolutionary and adaptive processes underlying cellular dynamics. When translating this biology into a stochastic system, it can be represented parsimoniously as the Markovian timeevolution (*96*) of the joint probability distribution *P*(*n, t*), which evolves according the *Chemical Master Equation* (*97*)

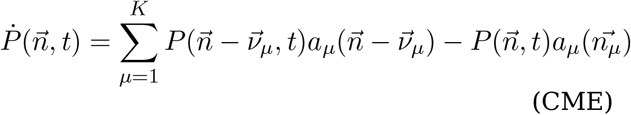

for *N* molecular species that participate in *K* chemical reactions, where 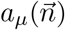 is the propensity of reaction *µ*. If 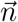 is a column vector of molecular numbers, then 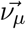 are the stoichiometric coefficients, i.e. *v*_*µ,i*_ = *v*_0;*µ,i*_ *− v*_1;*µ,i*_ is the coefficient of molecule *i* in reaction *µ*:

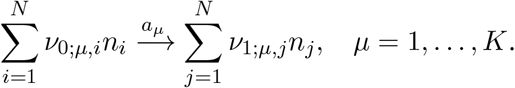

The CME captures exactly the stochastic dynamics of the considered chemical process. In this work, we model the coupled biochemical processes of gene regulation, transcription, and translation using a CME. Here, the considered species are transcripts (both pre-mRNAs and mRNAs) and proteins, and the chemical reactions are transcription (both initiation and elongation), translation, transcription factor binding and unbinding, splicing, proteasomal degradation, and RNA degradation. While analytic solutions are possible for linear systems, such as reaction networks composed solely of monomolecular (birth, death, and direct conversion *S*_*i*_ *→ S*_*j*_) reactions (*98*), they are generally not available for even simple non-linear systems. However, estimates of mean molecular numbers can be approximated using moment generating functions (*30*).

#### Sample-path simulation

Stochastic trajectories for the dynamics defined by the CME can be sampled exactly (i.e. simulated computationally) by stochastic simulation algorithms (SSAs), commonly known as “Gillespie” algorithms (*99*). Given the state 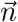, reaction *µ* fires in the next “infinitesimal” time interval [*t, t* +*dt*], with probability 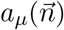, so that waiting times between events are exponentially distributed. The direct method iteratively draws the next event (which reaction fires next, and at what time), so that the computational cost per event grows with the number of reactions (*99*). Rejection-based SSA with composition-rejection search (RSSA-CR) (*64*) is an SSA variant which amortizes propensity computations, resulting in cost per event that is nearly independent of network size.

Simulating every event is expensive for largescale biological systems with fast reactions. This motivated the development of approximate methods, known as tau-leaping (*65*), which advance time in steps of duration *dt* = *τ* and fire each reaction Poisson 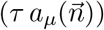 times, holding propensities fixed during the step. As such, the approximation only holds well when propensities don’t vary too much, and fails especially in the case of “critical” reactions involving lowcopy-number species, at which point some of the counts can even become negative. To mitigate this, many augmentations to the original algorithm have been proposed, trading off speed against accuracy. Negative counts can be prevented by clamping the counts to zero, or more accurately by repeatedly halving the *τ* -step using Poisson thinning, until no counts remain negative. Alternatively, the “critical” reactions, whose reactants are within *n*_*c*_ firings of exhaustion, can be simulated using an exact SSA algorithm, a procedure known as Critical tau-leaping (*69*). The *τ* value can also be chosen adaptively, so that the expected relative change in propensities during a step is bounded by an *ϵ* (*100*). Recently, an exact tau-leaping algorithm has been proposed, termed TauSplitting, which recursively bisects the time interval to be simulated until every propensity is “stable”, sampling the correct CME distribution without having to step through each event (*101*). JumpProcesses.jl (*34*) supplies many of these algorithms, and we are in the process of incorporating the rest into the library.

### B. A concrete model of intra-cell gene regulation

#### B.1 Transcriptional dynamics

Complex dynamics in our simulations arise from the effective interactions between molecular species. In particular, we are interested in the dynamics of DNA binding proteins, such as transcription factors, activators, repressors, enhancers, and silencers, that together regulate eukaryotic gene transcription (*102*). Our model is inspired by previous biophysical models of promoter regulation which have led to our current understanding of the role of promoter architecture in transcriptional dynamics (*13* , *23* –*25*). The foundation of our transcriptional dynamics simulation approach is the holo-promoter of each gene, which consists of its canonical promoter element and the collection of transcriptional activators and repressors which are associated with that gene (Fig. 1). We use the term holopromoter to include distant enhancer or silencer elements that may influence transcriptional dynamics from outside the canonical promoter or promoter-proximal regions.

Instead of assuming a quasi-steady promoter configuration, which would be adequate for reproducing some features of gene expression dynamics—in particular fold changes (*24*)—we have instead chosen to lift the time-scale separation up to the binding reactions of the individual transcription factors, such that the resulting changes in promoter configuration are in aggregate still represented explicitly as state. This not only increases the expressivity of this model as compared to methods that apply quasiequilibrium assumptions or time-scale separations at later stages of the gene expression cascade or even represent it as a single function in its entirety, but also makes this model more generally applicable, albeit at the cost of computational complexity and simulation time. We feel that this trade off is worthwhile, because important features of gene expression, such as mean-expression-independent variances or transcriptional bursting, can be easily captured in this framework.

Other simulation methods, such as the single-cell expression simulation software dyngen (*67*), use Hill-type propensity functions with high hill coefficients to model transcriptional regulation. It is usually assumed that this form of the propensity function is valid in the quasi-equilibrium limit of fast-promoter switching but, even in this regime, Holehouse et al. have shown that such propensity functions give incorrect protein number distributions, and can lead to incorrect predictions for the switching properties of genetic feedback circuits (*70*). In our model, while Hill-type propensity functions with large Hill coefficients are not assumed for the transcriptional propensities, they can nevertheless be realized in multiple ways, for example, by assuming a multimeric activation or repression function, or by combining multiple activators or repressors with cooperative binding for a given holo-promoter.

#### B.2 Canonical promoter

The canonical promoter models the binding of RNA polymerase and other proteins which are required to begin transcription. In our model, the promoter can exist in either an active or inactive state, which is explicitly modeled in the stochastic simulation. The promoter is assumed to enter the active state from the inactive state with a rate 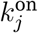, and switches back at a rate 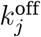. At equilibrium, the probability of being in the on state would be simply:

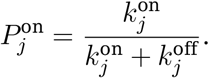

We model the transitions explicitly, however, as using the equilibrium fractional activity makes it difficult to encode the variance of mRNA independently from its mean. The mean mRNA levels for this model at equilibrium are proportional to the fractional activation 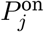, however, the variance is a function of the mean mRNA expression and of 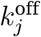 (*25*), which cannot be varied independently.

#### B.3 “holo”-promoter

The rate of transcription depends on the kinetics of the canonical promoter’s transitions between the active and inactive states. These kinetics depend on three factors, the “on-rate”, “off-rate”, and the concentration of RNAP as described above. In this model, the transcription factors associated with each holo-promoter act to adjust the rates 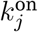 and 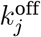. Transcription factors (TFs) thus either act as transcriptional activators, or transcriptional repressors. These classes include the traditional prokaryote inspired promoter-adjacent activators and repressors, as well as eukaryotic enhancers and silencers which may be distant to a given genes transcriptional start site. In general, a system may include *j* genes, and *i* TF binding sites, *i* ≠ *j*. Thus each gene may have zero, one, or many activators and repressors associated with its holo-promoter. We choose to express the combined activity of the activators *β*_ACT_ and repressors *β*_REP_ as the generalized mean of their individual fractional activations so that for *R ∈ {*ACT, REP*}*,

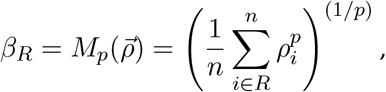

explicitly including the special case

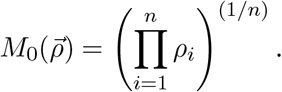

Thus, the use of the generalized mean permits as special cases the harmonic mean *M*_*−*1_, the geometric mean *M*_0_, the arithmetic mean *M*_1_, the quadratic mean *M*_2_, and as limiting cases also the minimum *M*_*−∞*_ and maximum *M*_*∞*_. Then the activation and inactivation rates for each gene 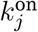 and 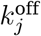 are given as a product of the unregulated on and off rates 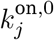 and 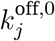 and the aggregation functions *β*_*R*_. Thus 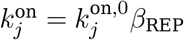 and 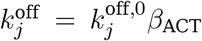. Thus the action of activators and repressors combine to collectively affect the activation and inactivation rates of the transcriptional initiation complex. These rates determine the average probability of finding the initiation complex in the active state as well as the rate of transitions back and forth from the active to inactive states. For the minimum or geometric mean aggregation functions, when any single repressor is fully bound, *β*_REP_ *→* 0, and the on-rate goes to 0. Alternatively, only when every activator is fully bound, *β*_ACT_ *→* 0 and the off-rate goes to zero. Here our model differs from other transcriptional models of activation, e.g. (*25*), which model activators as adjusting the rates of transcription by a multiplicative factor, however our model is in accord with experiments which show that activators alter the propensity of transcription initiation, but not the duration of each transcription event (*103*). Nevertheless, activators which alter the rate of triggering an elongation are interesting extensions to our model which we aim to include in future versions.

#### B.4. Elongation complexes

When in the active state (as described above), the promoter of gene *j* will trigger an elongation complex at a rate 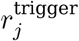 [RNAP]. Elongation complexes then form pre-mRNAs at a rate 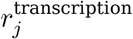 per complex which can be unique for each gene *j* and is in general a function of the pre-mRNA length. Elongation complexes can also abort transcription at a rate *λ*^elongation^ per complex. Completion or abortion of transcription releases the RNA polymerase back into the pool of free polymerases. The inclusion of the elongation complex in the model allows the transcription rate to implicitly be a function of both the triggering rate and the length of the unspliced transcript, longer transcripts will take longer to transcribe. However, because multiple elongation complexes can reside on a single transcribed gene (*104*), their production can still be triggered at a higher rate. This decoupling of transcriptional initiation and completion allows for the implicit inclusion of time-delays in the model, and may point to a quantifiable role for introns in the dynamics of eukaryotic transcription.

#### B.5. Processing

In eukaryotic organisms, splicing contributes to the overall dynamics of gene expression, and we model it explicitly. We assume that spliceosome availability is not the limiting factor for pre-mRNA processing, and that splicing is rate-limited by the availability of pre-mRNAs. This is supported by evidence that most splicing occurs co-transcriptionally and is surprisingly fast (*105*) and that while important, splicing delays seem to be signal-dependant rather than a passive effect of spliceosome availability (*106*).

In our model, each pre-mRNA is assigned its own 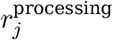 rate, which implicitly accounts for various factors that may affect pre-mRNA processing (*107*), such as splicing, capping, end-processing, polyadenylation, and signaling based delays, as well as spliceosome availability. While not modeled explicitly, adjusting 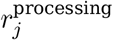 allows us to control the splicing rates of each pre-mRNA individually. In some cases, such as nutrient limited growth, it may be appropriate to include overall spliceosome availability, or to include a more mechanistically focused model of the splicing process; such model variants can be configured through overlay reactions, then optionally setting 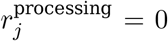. Transcripts are degraded with distinct rates 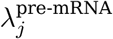 and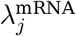.

#### B.6. Translation

Protein concentration is a function of protein production and degradation which are both modeled stochastically. Production is proportional to the concentration of the corresponding mRNA species. In addition, protein concentrations are controlled by two global factors which model the availability of ribosomes for translation and of proteasomes for degradation and which affect the production 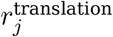, and degradation 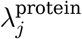, respectively for every gene in the simulation. For each gene *j* these rates are also a function of distinct per-gene translation and degradation rates, 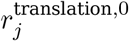, And 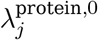, thus:

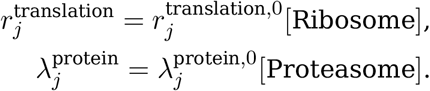

The ribosome and proteasome can be included in the stochastic simulation as separate genes. When this approach is used, we assume that the ribosome and proteasome are single genes which could represent, for example, them being the limiting component of each complex. For simplicity, these need not be simulated explicitly, and can instead be included simply as molecular species (which may be exogenously controlled). These simplifications are not required however, and the reaction overlay system can be used to model those molecular species as multi-gene complexes explicitly, including e.g. the rRNA and ribosmomal protein genes.

#### B.7. Multifate and proteolytic extension

The multi-fate system is a transcriptional regulation system designed (*55*) to have multiple stable states of transcription factor expression depending on the starting conditions and the degradation rates of the transcription factors. *GeneRegulatorySystems.jl* can be used to simulate these multistable gene networks, as demonstrated with a grid search of starting concentrations of the two monomer transcription factors (Fig. S1). In the original multi-fate design paper, it was noted that changing the rate of protein degradation could drive the system from the region of mono-stability to one of bistability. Taking this as inspiration, we include mutual proteolytic repression directly, so the system naturally undergoes this transition as the controlling timing factor degrades.

### C. Moment closure approximations for a single gene cascade. These equations were analytically derived, but in the context of a reified *GeneRegulatorySystems.jl* model are alternatively available automatically via MomentClosure.jl

#### Promoter state

The expectation values for the fraction of time the promoter is in the active state (*A*) and inactive state (*I*) are

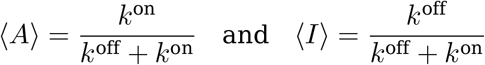

and the second moments are equal to the first moments:

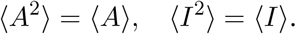

**Figure S1:**
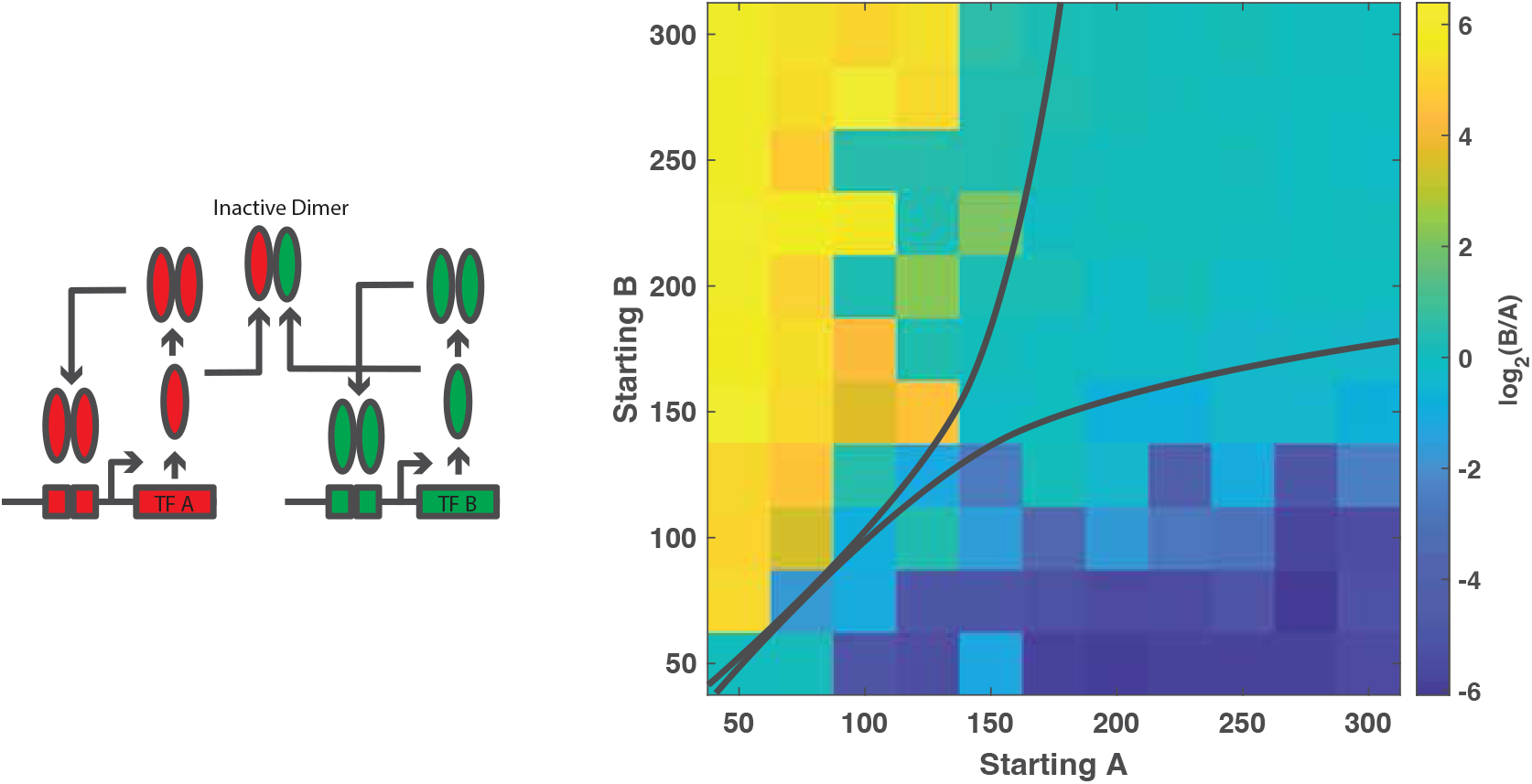
A grid search over starting concentrations of multifate transcription factors A and B demonstrating multistability.

As this is a binomial process, if *τ* = ⟨*A*⟩, its variance is given by Var(*A*) = *τ*(1 *− τ*).

#### Elongation Complexes

For elongation complexes, we have the following analytic expressions for steady state first and second moments

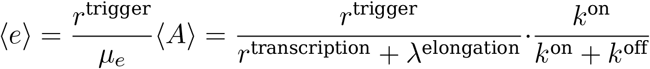

And

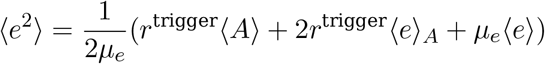

where

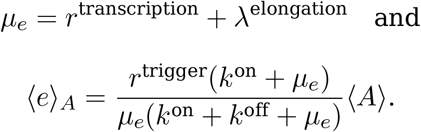

Therefore, the full expression for the second moment for elongation complexes is given by:

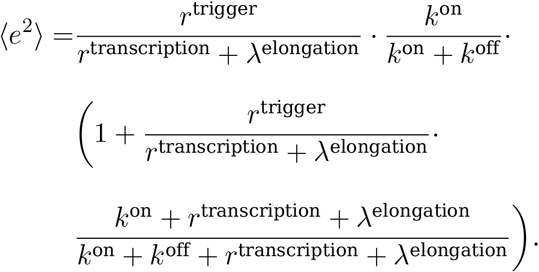

#### Nascent (pre)-mRNA

We can use the same approach to calculate the next moments for premRNAs *n*. The mean value is easier to calculate, as it is simply the mean value of the proceeding species multiplied by the ratio of all production to all decay constants

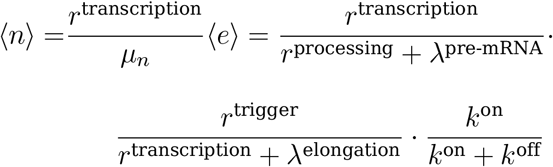

With

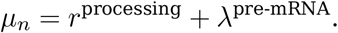

The second moment can also be calculated as

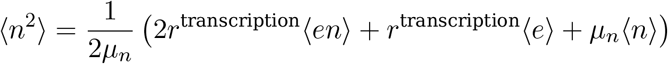

which includes a mixed moment term ⟨*en*⟩ given by

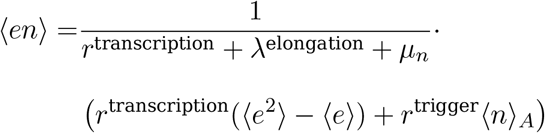

and requiring the term

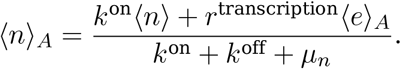

**Mature mRNAs** The first moment of the mRNA distribution is given as

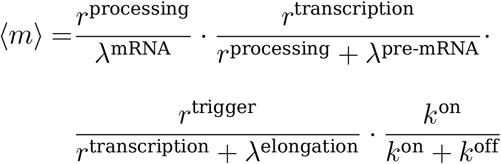

and for the second moment we have a similar expression for mRNA as we had for pre-mRNA,

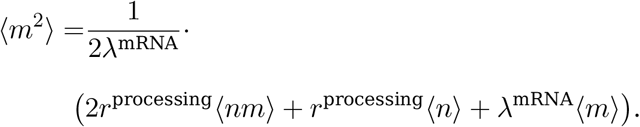

Now we have additional mixed moments to include, namely:

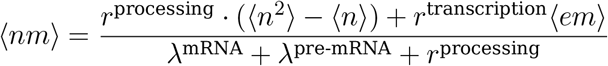

And

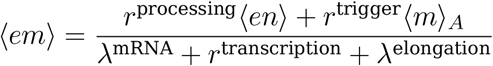

With

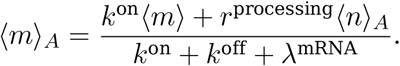

**Proteins** Finally at the protein level we have the first moment given by

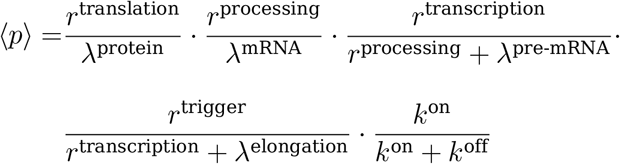

and the second moment given by

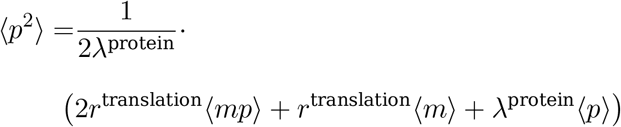

requiring the following mixed moments to calculate:

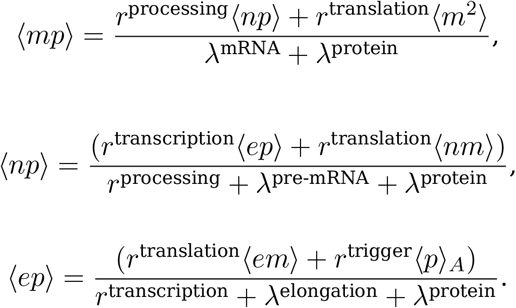

Similar derivations can be found in (*108*).

### D. Example: The *C. elegans* endomesoderm network

We show how *GeneRegulatorySystems.jl* can encode a published experimental design and compare its output directly to measured data. As an example, we use a well characterized developmental GRN, the *C. elegans* endomesoderm network, which directs daughters of the EMS blastomere, E and MS, to adopt endodermal and mesodermal fates, respectively. This specification is established through a mixture of exogenous factors (signaling from neighboring P2 cells), and endogenous programming ensuring zygotic transcription factors are expressed in sequence and with the right timing (*109*). The *C. elegans* embryo makes for an ideal case study due to its stereotyped early development (*110*), reproducible cell cycle lengths (*111*) and well-annotated single-cell omics data (*112*), facilitating direct comparison of simulated measurements with experimental data, without the need for pseudotime inference.

We combined two complementary datasets into a single schedule (Fig. S2A): single-nucleus RNA sequencing (snRNA-seq) profiles of wildtype embryos across the early E and MS lineages from van der Burght et al. (*112*), and endogenously tagged fluorescence reporters of key transcription factors involved in the E lineage differentiation under gene knockouts from Ewe et al. (*109*). For the schedule timing, we derived stage durations from lineage-resolved imaging at 20^*°*^*C* (*111*), rescaled to 18^*°*^*C*, the growth temperature of the snRNA-seq embryos, via Arrhenius’ law with an apparent activation energy of 75 kJ/mol, so that every stage lasts 24% longer (*113* , *114*). As the snRNA-seq annotation splits each cell cycle into an early and late stage, we assigned the first third of each cycle to the early stage. Each simulated embryo starts as an EMS cell loaded with SKN-1, POP-1 and REF-1 protein, and branches at the EMS division into an MS and E lineage (which share their common EMS history). Signaling from P2, which in vivo involves the Wnt, MAPK and Src pathways (*109*), is represented by a single proxy molecular species, P2-signal. For comparison with the snRNAseq data, pre-mRNA counts are sampled every 50 s through a binomial extraction model with 20% capture efficiency (with 20 replicate embryos simulated within the same schedule). For the knockout experiments, the E lineage is simulated under ten measured genotypes (80 replicates each): genetic knockouts remove both promoter copies of the target gene, whereas RNAi knockdowns introduce an siRNA species that degrades the target RNA through an overlay reaction. Protein counts are read out through the same extraction model.

All simulated conditions share the same regulatory network and parameters (Fig. S2B). The E lineage topology follows Ewe et al. (*109*): a feedforward cascade of partially redundant GATA factors (MED-2/1, END-3/1, ELT-7/2), initiated by maternal SKN-1. We extended it with the MS branch, built around the two key MS regulators TBX-35 and CEH-51 (*115*). The two fates are differentiated by the P2-signal, which converts POP-1 from a repressor of the *end-3/1* genes into an activator (*109*) and represses *tbx-35* in E via the REF-1 family (*116*). Base rates are shared across genes and taken from measurements in *C. elegans* where available, such as transcription initiation (*117*) and mRNA decay (*118*), and from other systems otherwise (*119* –*121*) (Table S1). Ideally, these rates should also be adjusted depending on temperature like above, but for simplicity, here they were just selected from the higher ends of the intervals given in the preceding references. We assume a uniform elongation speed of 40 nt/s for all genes, as measured in *Drosophila* embryos (*122*), so that transcription time scales with gene length (taken from Ensembl, WBcel235 assembly, including introns (*123*)). Decay rates for mRNA and proteins follow the shared defaults with a few classes, distinguishing rapidly clearing early factors (SKN-1, MED-2/1) from stable late regulators. Regulatory edge Hill parameters were tuned by hand against the data.

The simulations reproduce the lineage- and stage-specific expression pattern of the snRNA-seq data, with mismatches mainly in the timing of expression onset and cessation (Fig. S2C, D). Under knockout, the model captures the partial redundancy of MED-2/1. Further down the cascade, the knockouts reveal negative cross-regulation among END-1, ELT-7 and ELT-2, which the model is able to express only partly (Fig. S2E). In particular, it misses the rise of ELT-2 in *end-1* and *elt-7* (RNAi) embryos, a limitation shared by the original ODE model used in Ewe et al. (*109*). Adding repression of *elt-2* by END-1 or ELT-7 did not resolve this: it produced the increase in the mutants only at the expense of the wildtype ELT-2 levels. This suggests that the interaction between these genes is more subtle and context-dependent than the current *GeneRegulatorySystems.jl* promoter model can accommodate, for example through chromatin state or other combinatorial effects. In fact, chromatin accessibility was profiled in the same nuclei as the snRNA-seq data (*112*), but we did not use it, as it would require an explicit model of chromatin state.

Our aim here was to present a minimal example of a schedule encoding an experimental design matched to data from multiple sources and modalities. Natural extensions to the model include cell volume and division, more complex promoter regulation, and a more detailed signaling network. Most of these, however, would add parameters to a model that is already arguably over-parameterized for the available data. Parameter inference with uncertainty quantification would reveal which interactions the data actually identify, and could be used to inform new experiments (*124*). This will be the subject of future work.

**Figure S2:**
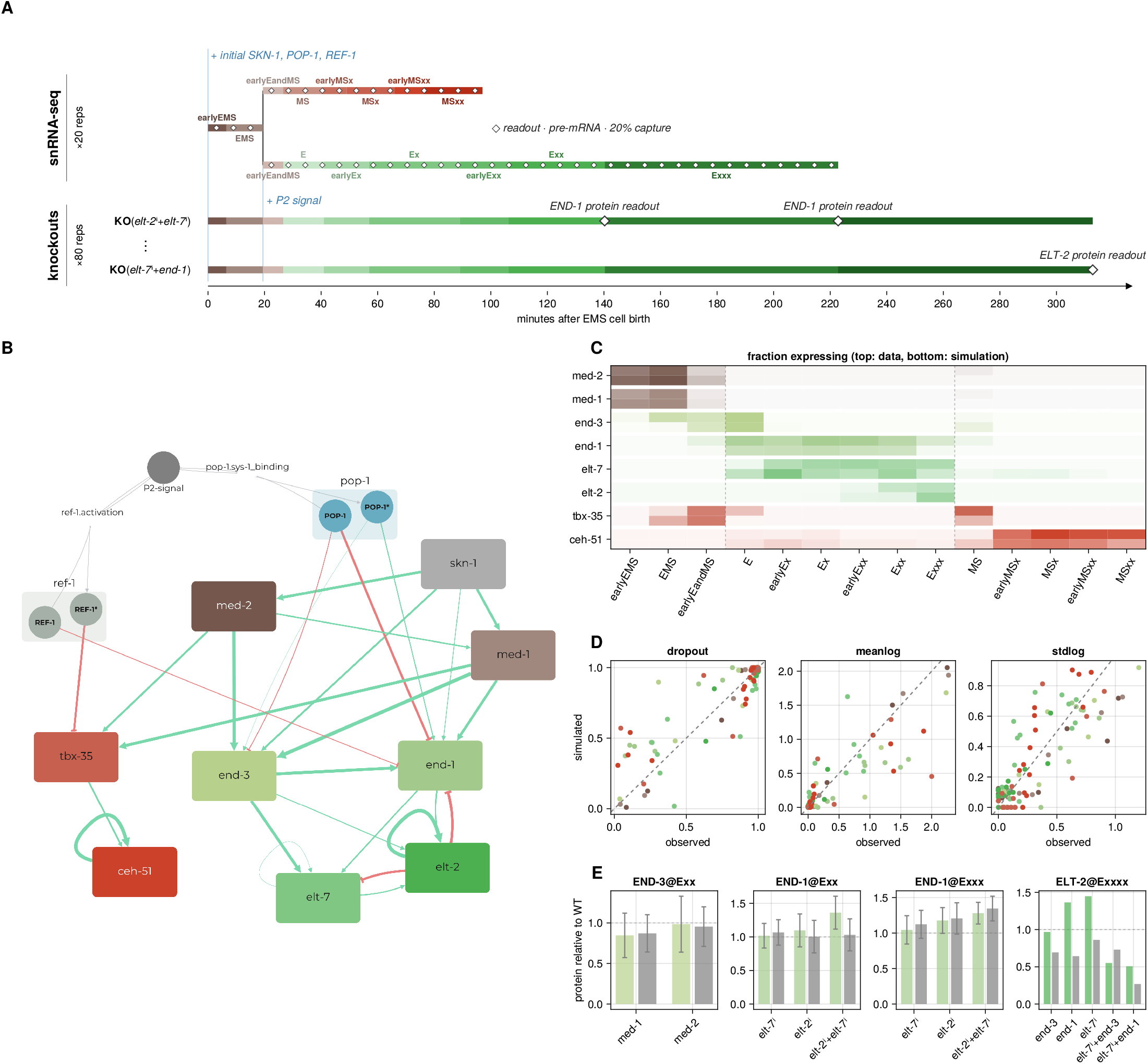
A single schedule integrates multimodal experimental data of *C. elegans* embryo development. A *GeneRegulatorySystems.jl* schedule was constructed to jointly express two experimental designs: wildtype cell differentiation with fixed stage durations and simulation branching, matched against snRNA-seq data, and knockout experiments under several genotypes (2/10 shown) **(A)**. A single regulatory network is shared across all branches. The regulatory edges, weighted by strength, and selected overlay reactions are shown **(B)**. The simulated transcriptome measurements were compared against the snRNA-seq data (*112*) as the fraction of nuclei with nonzero counts per gene and stage **(C)**, and as per-stage dropout, mean and standard deviation of log counts (colored by gene as in B) **(D)**. Simulated protein levels under knockout or RNAi knockdown (gray), relative to wildtype levels (dashed line), were compared against reporter fluorescence measurements (green) (*109*) **(E)**. Error bars reflect one standard deviation.

**Table S1:** Reaction base rate constants. These rates are used for genes of the given types in the presented examples and are collected here for convenience; their actual specification can be found in the respective simulation schedules. Rates marked ⊛ will be automatically adjusted (Methods K). The *C. elegans* column shows the default gene rates, which get partially overridden for certain genes. L stands for gene length in nucleotides. For details, see https://github.com/drostlab/grs-paper.

| **Reaction** | **Rate constant** [s<sup>−1</sup>] | **Default** | **Differentiator** | **Timer** | **Benchmark** | **C. elegans** |
|---|---|---|---|---|---|---|
| activation | *k*<sup>on,0</sup> | 2.5 | 2.5 | 0.1 | 0.0003 | 0.015 |
| deactivation | *k*<sup>off,0</sup> | 10.0 | 10.0 | 0.1 | 0.002 | 10.0 |
| initiation | *r*<sup>trigger</sup> | 6.6 · 10<sup>−7</sup> | 6.6 · 10<sup>−7</sup> | 6.6 · 10<sup>−7</sup> | 1.94 · 10<sup>−7</sup> | 4 · 10<sup>−7</sup> |
| abortion | λ<sup>elongation</sup> | 0.01 | 0.01 | 0.01 | 0.01 | 4 · 10<sup>−4</sup> |
| transcription | *r*<sup>transcription</sup> | 0.001 | 0.015 | 2.5 · 10<sup>−7</sup> ⊛ | 0.001 | 40/*L* |
| pre-mRNA decay | λ<sup>pre-mRNA</sup> | 0.001 | 0.001 | 0.001 | 0.001 | 0.001 |
| processing | *r*<sup>processing</sup> | 0.02 | 0.02 | 0.02 | 0.02 | 8.3 · 10<sup>−3</sup> |
| mRNA decay | λ<sup>mRNA</sup> | 0.001 | 0.0075 | 0.01 | 6.4 · 10<sup>−5</sup> | 5.8 · 10<sup>−4</sup> |
| translation | *r*<sup>translation,0</sup> | 2.5 · 10<sup>−9</sup> | 2.5 · 10<sup>−9</sup> | 10<sup>−9</sup> | 7.2 · 10<sup>−9</sup> | 3 · 10<sup>−8</sup> |
| protein decay | λ<sup>protein,0</sup> | 3 · 10<sup>−10</sup> | 2 · 10<sup>−10</sup> | 1.0 ⊛ | 2.4 · 10<sup>−11</sup> | 5.8 · 10<sup>−10</sup> |

**Table S2:** Simulation cost on the 10,000-gene bifurcating-cycle model used in dyngen’s runtime benchmark (*67*). The model generated by dyngen (modeled timespan of 552 h) was imported into *GeneRegulatorySystems.jl*, reproducing dyngen’s quasi-steady-state promoter architecture, so that both engines simulate identical chemical reaction networks. Build time is the time taken to compile the model into a simulator from a cold start; dyngen performs no caching, so it pays this cost for every network instance it generates. Runtime is single-threaded wall-clock per trajectory, averaged over three trajectories. Fixed-step methods use *τ* = (300*/*3600) s^*−*1^ = (5 min)^*−*1^. dyngen’s tau-leaping clamps species that would become negative to zero, an additional source of bias; we compare against more accurate tau leaping variants (App. A).

| **Engine** | **Build (s)** | **Method** | **Runtime (s/traj.)** |
|---|---|---|---|
| dyngen | 320.7 | tau-leaping, fixed τ, clamped negatives | 26.7 |
| *GeneRegulatorySystems.jl* | 88.4 | tau-leaping, fixed τ, clamped negatives | 40.7 |
|  |  | tau-leaping, fixed τ, thinned negatives | 67.2 |
|  |  | adaptive tau-leaping, ϵ = 0.05 | 111.8 |
|  |  | hybrid, fixed τ, threshold *n*<sub>*c*</sub> = 10 | 425.9 |
|  |  | exact SSA (TauSplitting) | 884.4 |

**Figure S3:**
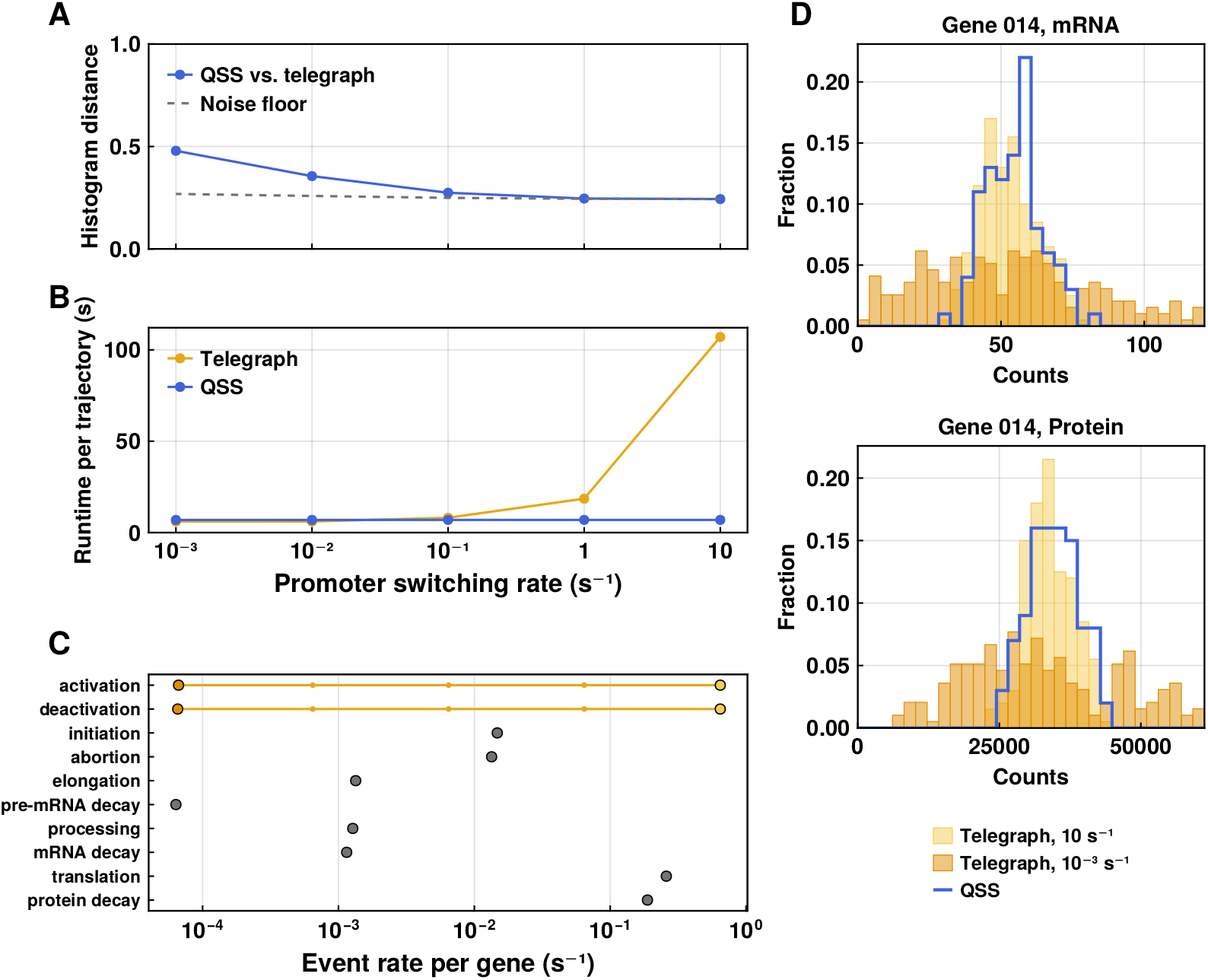
Comparing the explicit promoter switching (telegraph) model to the quasi-steadystate (QSS) approximation. For each promoter model, a Kronecker network of 512 genes was simulated for 172,800 s of model time and regularly sampled at ten time points. The telegraph model switches each promoter with activation and deactivation of *d/*2, for switching rate *d* from 10^*−*3^ to 10 s^*−*1^. The QSS model replaces the promoter with its steady-state activity (making it independent of *d*). **(A)** Histogram distance between QSS and telegraph model (*N* = 100 trajectories, with the sampling noise floor dashed, as in Methods M.6). **(B)** Runtime per trajectory. **(C)** Measured event rates per gene for each reaction kind. Effective promoter activation and deactivation is shown for different switching rates. **(D)** mRNA and protein counts histograms for one example gene at steadystate (*t* = 172, 800), comparing fast and slow switching versus the QSS model.

